# A comprehensive comparison on clustering methods for multi-slide spatially resolved transcriptomics data analysis

**DOI:** 10.1101/2025.01.19.633631

**Authors:** Caiwei Xiong, Huang Shuai, Muqing Zhou, Yiyan Zhang, Wenrong Wu, Xihao Li, Huaxiu Yao, Jiawen Chen, Yun Li

**Affiliations:** Department of Biostatistics, UNC-Chapel Hill; Department of Genetics, UNC-Chapel Hill; Department of Computer science, UNC-Chapel Hill

**Keywords:** spatial transcriptomics, clustering, multi-slide clustering, evaluation

## Abstract

Spatial transcriptomics (ST) data, by providing spatial information, enables simultaneous analysis of gene expression distributions and their spatial patterns within tissue. Clustering or spatial domain detection represents an essential methodology for ST data, facilitating the exploration of spatial organizations with shared gene expression or histological characteristics. Traditionally, clustering algorithms for ST have focused on individual tissue sections. However, the emergence of numerous contiguous tissue sections derived from the same or similar tissue specimens within or across individuals has led to the development of multi-slide clustering methods. In this study, we assess seven single-slide and three multi-slide clustering methods on two simulated datasets and three real datasets. Additionally, we investigate the effectiveness of pre-processing techniques, including spatial coordinate alignment (for example, PASTE) and gene expression batch effect removal (for example, Harmony), on clustering performance. Our study provides a comprehensive comparison of clustering methods for multi-slide ST data, serving as a practical guide for method selection in various scenarios.

## 1 Introduction

The intricate interaction of gene expression across cells is fundamental to our understanding of biological systems. The advent of ST technologies marks a significant milestone in this exploration, offering a window into gene expression landscapes within their spatial context in tissue. ST technologies mainly fall into two categories, nextgeneration sequencing (NGS)-based approaches and imaging-based approaches [1] [2]. NGS-based approaches involve the collection of RNA on microarray slides embedded with spatial barcodes prior to the reverse transcription step, guaranteeing that every transcript could be linked to its initial location through its unique positional molecular barcode [3]. NGS-based approaches encompass technologies including 10X Genomics Visium [4], Slide-seq [5], as well as more recent technologies like deterministic barcoding in tissue for spatial omics sequencing (DBiT-seq)[6], spatio-temporal enhanced resolution omics sequencing (Stero-seq) [7], and polony (or DNA cluster)indexed library sequencing (PIXEL-seq) [8]. Imaging-based approaches can be broadly categorized into two groups: situ sequencing (ISS) and in situ hybridization (ISH). Within the ISS category, RNA undergoes reverse transcription, followed by amplification through rolling circle amplification, and finally sequencing [9]. Spatially resolved transcript amplicon readout mapping (STARMap) [10], fluorescent ISS (FISSEQ) [11] and expansion sequencing (ExSeq) [12] follow this ISS approach. In contrast, ISH technologies like multiplexed error-robust fluorescence ISH (MERFISH) [13] allow several sequential circles of hybridization and imaging combined with barcoding.

ST has become a cornerstone in various biological research fields [14]. For example, ST enables detailed mapping of cell types for diverse tissue types, encompassing brain [15] [16] [17], heart [18], and lung [19]. Additionally, ST is instrumental in illustrating cellular interactions within spatial contexts, with studies covering areas such as mouse visual cortex [10], Alzheimer’s disease [20], and brain injury [21]. With these two applications combined, ST has the capacity to elucidate how different cell types interplay with one another across spatial locations and help reveal molecular interaction among tissue constituents. Such interaction includes tumors and their neighboring environment [22], the infiltration of immune cells within tissues, and the formation of developmental gradients [23]. Despite these advancements, an essential initial step in analyzing ST data involves using clustering algorithms to identify spatial domains within tissues. Investigating these domains and their associated genes offers insights into tissue organization and disease progression, highlighting the multifaceted applications and significance of ST data in biological research.

Clustering algorithm categorizes groups of spots/cells with similar intrinsic characteristics. Compared with the single cell RNA-seq(scRNA-seq) clustering methods, ST clustering methods incorporate spatial information, which has been proven to improve clustering performance in multiple studies [14] [24] [25]. The premise behind is that spots in close proximity are more likely to share similar characteristics. Thus, the key to effective ST clustering is the strategic use of spatial information. For instance, graph convolutional network (GCN)-based approaches use spatial locations to construct graphs, representing spatial neighborhoods. Spatial information can also be derived from similarity in histology images or cell-type proportions. SpaGCN, for example, integrates histology images, while STAGATE and BASS account for celltype proportions, contributing to a deeper and more interpretable understanding of spatial domains.

As ST technology gains traction, researchers are producing increasing amounts of data, often encompassing multiple contiguous tissue sections from the same sample or comparable tissue specimens from different individuals. This surge in data generation has created a need for methods capable of analyzing multiple slides simultaneously. Similar to scRNA-seq analysis, analyzing multiple datasets requires data preprocessing techniques like harmonization or batch effect removal to ensure consistency across samples. Given the inherent spatial aspect of ST data, aligning spatial locations across different slides is also crucial. Methods like PASTE and STalign have been developed to align spatial locations of spots across multiple slides, ensuring that spots from similar biological structures occupy consistent spatial coordinates. These preprocessing techniques form the foundation for clustering methods designed for multi-slide ST analysis, such as BASS. While spatial domain detection methods for single-slide clustering have been extensively evaluated [26], there has been limited research comprehensively assessing both single-slide and multi-slide clustering approaches. Key questions remain: does analyzing multiple slides simultaneously improve clustering performance? What impact does data preprocessing have on multi-slide clustering? Can single-slide methods, with spatial alignment and harmonization, outperform multi-slide methods? Answering these questions could provide valuable insights into optimizing ST analysis for larger and more complex datasets.

In this review, we provide a comprehensive summary and comparison of clustering methods for ST data. We outline ten state-of-art ST clustering methods, including seven single-slide and three multi-slide methods, emphasizing crucial aspects such as the statistical or machine learning methodology employed, and unique features specific to each method. We assess the performance of these methods using two simulated and three real ST datasets. Additionally, we evaluate the effectiveness of data pre-processing methods, including Harmony for gene expression harmonization and PASTE for spatial coordinate alignment, for multi-slide data. Lastly, we offer practical guidelines and underscore the advantages and limitations of these methods when applied to single- and multi-slide ST data.

## 2 Clustering methods for ST data

In recent years, researchers have proposed a diverse array of ST clustering methods. Existing clustering approaches for ST data can be broadly categorized into three main groups (Fig 1): Bayesian methods, graph network based methods, and other methods. Bayesian modeling methods assume that gene expression data follow a specific distribution conditional on cluster/layer and/or spatial coordinates. These methods estimate the distribution parameters from ST data, thereby acquiring the ability to predict clusters. Graph network methods utilize graph structures as encoders or feature selection tools to facilitate clustering or feature selection. These methods employ contrastive learning or minimize reconstruction loss during the training phase, enabling clustering upon model completion. Other methods, such as Seurat, Stardust, and stLearn, leverage specially-designed integration of gene expression and spatial coordinates as input for clustering algorithms. In this work, we examine ten state-of-the-art methods: Seurat, stLearn, Stardust, SpaGCN, GraphST, BayesSpace, STAGATE, iSC.MEB, BASS, and MAPLE. Notably, iSC.MEB, BASS, and MAPLE are specifically designed for multi-slide clustering. Table 1 provides a summary of main features of these methods. Next, we provide detailed methodological summarization for each method evaluated.

**Table 1.**
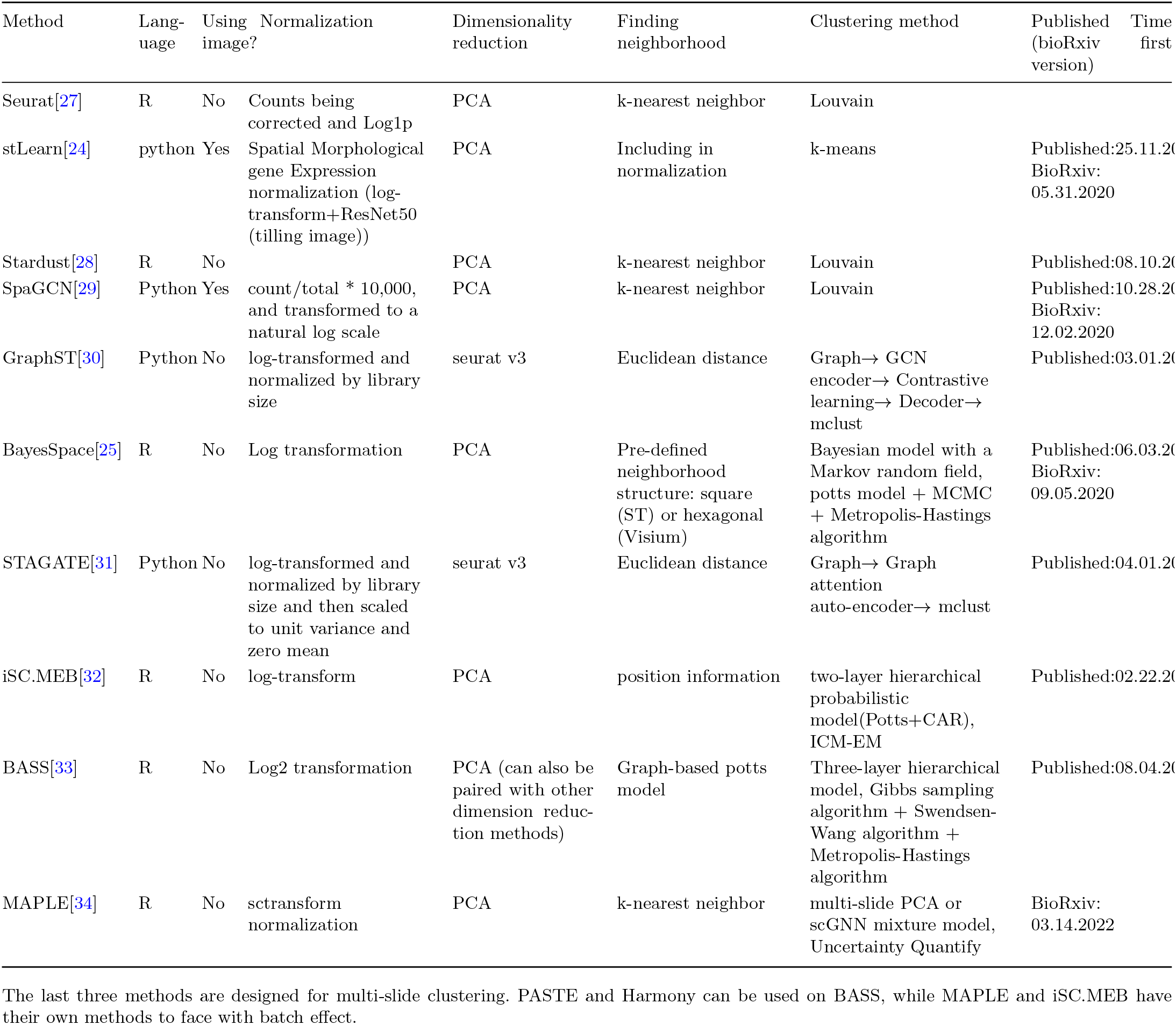
ST clustering methods overview.

**Fig. 1.**
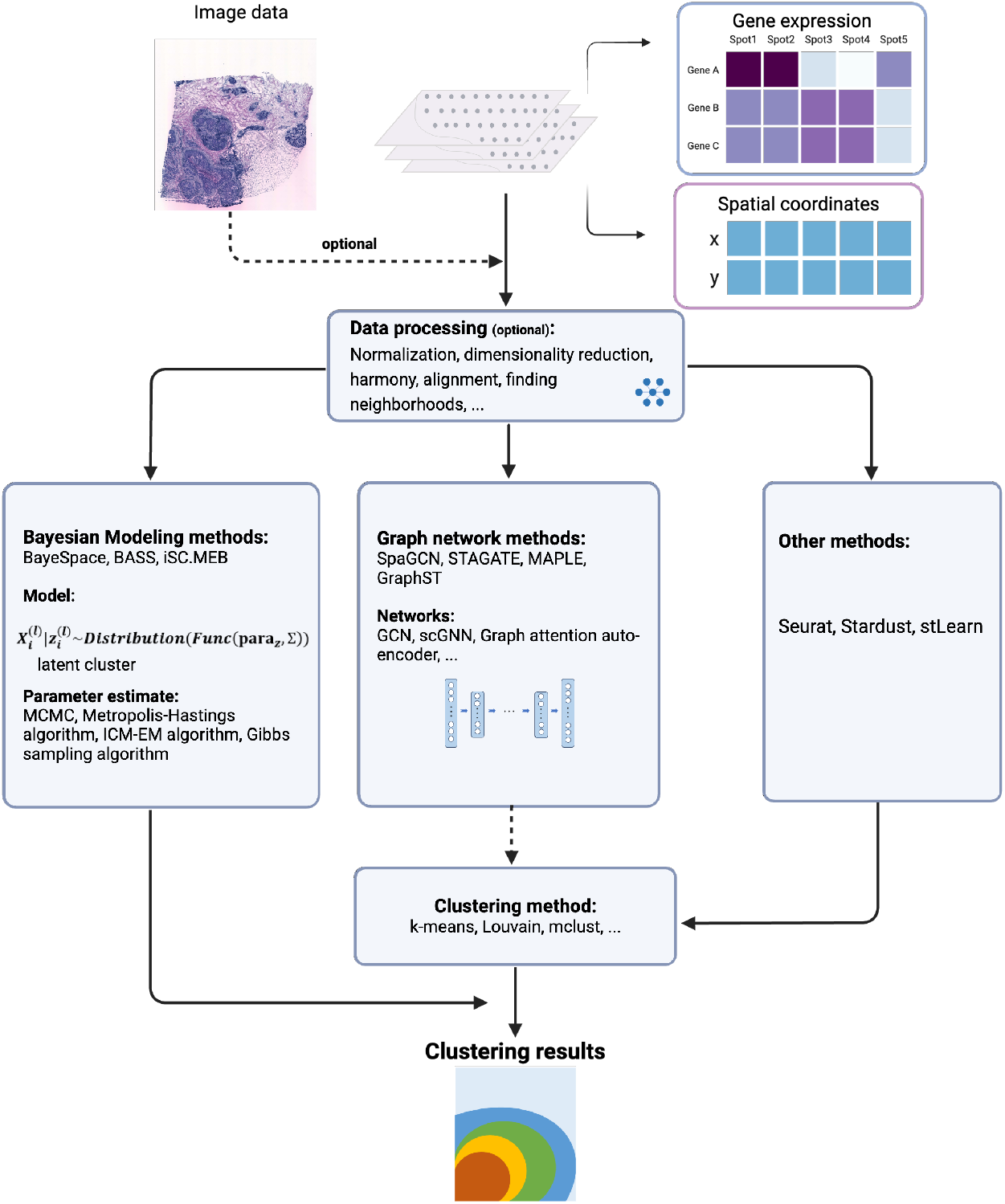
Summary of ST clustering methods. ST clustering methods take (pre-processed) gene expression data, spatial data and optional image data from multiple slides as input (top panel). The core algorithm of current ST clustering methods can be classified into three main categories: Bayesian methods, graph network methods and other methods (center panel). The output of ST clustering methods is a predicted cluster for each spot (bottom panel where each color represents a cluster). para: estimated or pre-specified parameters. Func: a function of parameters and clusters. X: the expression feature matrix. Z: spatial domain labels across all sections.

Seurat [27] is a comprehensive R toolkit equipped with functionalities for data pre-processing, dimension reduction and gene expression analysis. The default Seurat workflow for clustering ST data entails dimensionality reduction via principal component analysis (PCA), identifying neighboring cells using the k-nearest neighbor (k-NN) algorithm with the Euclidean distance metric, and clustering using the Louvain algorithm. Alternatively, various dimensionality reduction techniques can be applied, such as Independent Component Analysis (ICA), Supervised Latent Semantic Indexing (SLSI), Supervised Principal Component Analysis (SPCA), t-distributed Stochastic Neighbor Embedding (t-SNE), and Uniform Manifold Approximation and Projection (UMAP). In conjunction, different distance metrics, such as cosine, Manhattan, and Hamming, can be utilized for Approximate Nearest Neighbors (annoy). Additionally, a variety of clustering methods, including the Smart Local Moving (SLM) algorithm and the Leiden algorithm, can be employed to enhance performance and insights.

Stardust [28] offers a simple and flexible approach to incorporate both spatial and transcriptomic data by replacing the Euclidean distance matrix used in Seurat is replaced by a combination of two alternative matrices, one representing transcriptional data and the other indicating the spatial positions of the spots. This allows either manual or fully automated tuning of parameters. In the manual method, a parameter called “spaceWeight” specifies the influence of the spatial-based measure on the overall metric. The automatic approach computes the final distance matrix as a blend of both spatial and transcript information, considering both normalized and non-normalized expression distance distribution. The cluster method is also based on the Louvain algorithm, same as Seurat. In our analysis, we opt for the automatic distance matrix due to its more robust performance compared to manual parameter specification. stLearn [24] is a method that places significant emphasis on data preprocessing, amalgamating spatial location, tissue morphology from images, and gene expression profiles. The first step of stLearn is Spatial Morphological gene Expression normalization(SME normalization). The configuration of spot morphology involves capturing the top 50 features from spot image tiles through a pre-trained ResNet50 model. A measure of morphological similarity is computed by evaluating the cosine distance between feature vectors. For each gene within any particular spot, its expression value is computed as the average of morphological similarity-weighted expression values from its neighboring spots. The next step is spatial clustering (SMEclust) using the SME normalized matrix. It begins with PCA/UMAP dimensionality reduction, followed by the establishment of a k-nearest neighbor graph, and the implementation of global clustering methods such as Louvain or k-means clustering.

BASS (Bayesian Analytics for Spatial Segmentation)[33] operates within a Bayesian hierarchical framework to enable comprehensive analysis in multi-scale and multi-slide scenarios. For gene expression, BASS aggregates cells across all tissue section, conducts library size normalization followed by a log2-transformation, and performs dimension reduction on the normalized expression matrix. BASS uses spatial location data to construct a cell-to-cell neighborhood graph for each tissue section by identifying the k-nearest neighbors for every cell. In particular, BASS assumes that the expression distribution is conditional on cell type information, while the cell type follows a categorical distribution contingent on the underlying spatial domain. The spatial domain label is determined based on the neighborhood graph through a homogeneous Potts model. To estimate the model’s parameters, BASS adopts a Gibbs sampling algorithm in conjunction with a Metropolis-Hastings algorithm, and additionally employ the Swendsen-Wang algorithm to achieve faster convergence.

SpaGCN [29] is a graph convolutional network approach designed to identify spatial domains and spatially variable genes. In addition to using gene expression and spatial information, it also embeds color information from histology images by using a weighted sum of mean RGB values of 50*×*50 pixels around each spot. Then the Euclidean distance between spots is calculated from spatial coordinates and the image color. Gene expressions are normalized and a graph convolutional network is iteratively trained while simultaneously optimizing cluster centroids. For the loss function, SpaGCN uses Kullback–Leibler (KL) divergence, which emphasizes spots assigned with high confidence and normalizes the contribution of each centroid. This approach ensures that larger clusters do not skew the representation in the hidden feature space.

iSC.MEB (integrated spatial clustering with hidden Markov random field using empirical Bayes) [32] is tailored for conducting comprehensive clustering analysis that addresses batch effects on low-dimensional representations across multiple ST datasets. iSC.MEB first applies PCA to the combined log-transformed expression matrix, yielding the top principal components (PCs). For the clustering process, iSC.MEB uses a two-layer hierarchical probabilistic model, encompassing a conditional distribution of expression matrix given unknown labels and a Potts model that fosters spatial consistency. Furthermore, iSC.MEB incorporates a conditional autoregressive (CAR) model that captures spatial correlation induced by neighboring microenvironments imposed on the batch-related part. The iSC.MEB algorithm utilizes an iterated conditional modes algorithm (ICM) combined with EM framework to iteratively estimate parameters, alternating between label assignment and batch-related embedding updates, deriving the evidence lower bound (ELBO) in the E-step, and optimizing the smoothness parameter through grid search, with the optimal number of clusters determined via MBIC after convergence.

STAGATE [31] adopts a graph attention auto-encoder framework to identify spatial domains by learning low-dimensional latent embeddings via integrating spatial information and gene expression profiles. Specifically, STAGATE first constructs the spatial neighbor network (SNN) based on the relative spatial locations of spots, and optionally introduces cell type-aware SNN by pruning the SNN based on the preclustering using only gene expressions. Then STAGATE learns low-dimensional latent embeddings with both spatial information and gene expressions via a graph attention auto-encoder. STAGATE adopts an attention mechanism in the middle layer of the encoder and decoder. The objective of STAGATE is to minimize the reconstruction loss of normalized expressions. Finally, the latent embeddings are used to identify spatial domains with various clustering algorithms, such as mclust and Louvain. When the number of domains is known, the mclust clustering algorithm will be employed. The procedures described above profile gene expression patterns in the context of 2D tissue sections. To accommodate multiple samples, the 3D SNN based on 3D space is introduced, where SNN between adjacent sections is constructed based on the aligned coordinates and a pre-defined radius.

GraphST [30] is a graph self-supervised contrastive learning method. It combines graph neural networks with self-supervised contrastive learning to learn informative and discriminative spot representations. Specifically, it first creates a corrupted graph by randomly shuffling gene expression vectors across spots while keeping the adjacency matrix of the original graph unchanged. Then with the original and corrupted graphs G and G’ as inputs, the Graph neural network (GNN) based encoder first generates two corresponding representation matrices. The contrastive learning is modeled with the original and corrupted graphs as inputs in order to learn the spot representations more accurately and output a refined representation. Then this latent representation is fed into a decoder to be reversed back into the raw gene expression space. The model is trained by minimizing the self-reconstruction loss of gene expressions. Finally, the reconstructed spatial gene expression from the decoder is used with mclust to cluster the spots into different spatial domains. To make it applicable for multiple samples, a joint neighborhood graph for multiple slices is constructed in the same way as with a single slice, after applying PASTE to first align the coordinates of different slices.

MAPLE [34] is a hybrid framework that includes a GNN to extract informative features from multi-slide ST data, and a Bayesian finite mixture model to detect spatially informed cell spot sub-populations and compare cell spot sub-population composition between samples while adjusting for possible confounding factors in multislide experimental designs.

BayesSpace [25] is a fully Bayesian statistical method that uses information from spatial neighborhoods for clustering and resolution enhancement. BayesSpace uses preprocessed ST (transformed, normalized and PCA) data to construct a Bayesian hierarchical model to infer spatial clusters. Potts model prior is used to encourage neighboring spots to belong to the same cluster. To enhance clustering resolution, each spot is segmented into sub-spots and gene expression of sub-spots are considered as latent variables in the model.

In summary, numerous innovative ST clustering methods have been developed and tailored specifically for ST data. These methods have demonstrated their potential in both simulated and real datasets. While several studies have evaluated performance of methods designed for single-slide data, to the best of our knowledge, there is a lack of a comprehensive evaluation of the performance in multi-slide settings. In this study, we employ two simulated and three real ST datasets (Table 2), to systematically and objectively assess the performance of these methods in the context of multi-slide clustering. This comprehensive evaluation aims to provide unbiased comparisons and thus offer practical guidelines for the usage of these methods.

**Table 2.** Summary of real datasets.

| Data type | Sample Size | Using Images? | Tissue | ref. | Link |
| --- | --- | --- | --- | --- | --- |
| 10x | 12 | Yes | Human dorsolateral prefrontal cortex (DLPFC) | [15] | https://research.libd.org/spatialLIBD/ |
| ST data | 11 | Yes | HER2+ breast cancer | [35] | https://zenodo.org/records/4751624 |
| MERFISH | 5 | No | Mouse hypothalamus | [36] | https://datadryad.org/stash/dataset/doi:10.5061/dryad.8t8s248 |

## 3 Results

To evaluate the performance of the aforementioned ten methods, we employed three real datasets (Table 2) and two simulated datasets. For quantitative assessment, we calculated the Adjusted Rand index (ARI) (Technical details) between every slice of ground truth labels and the labels predicted by the clustering methods. Additionally, we compared the performance through visual evaluation of clustering results. To compare the performance between single- and multi-slide clustering methods, we applied all ten clustering methods in both single- and multi-slide scenarios. Furthermore, to compare the impact of data preprocessing, specifically via PASTE and Harmony, we conducted three sets of analyses: (1) multi+Harmony: multi-slide clustering after Harmony correction, (2) multi+PASTE: multi-slide clustering after PASTE alignment, and (3) multi+PASTE+Harmony: multi-slide clustering after applying both Harmony and PASTE. We opted not to implement these preprocessing steps for the multi-slide clustering approaches (Maple and iSC.MEB), positing that these methods inherently have their own strategies to aggregate data from multiple samples. However, for BASS, which employs Harmony as the default, we hypothesized that incorporating a technique to reduce batch effects could potentially improve its performance. In summary, we performed the analysis with Harmony and PASTE added using BASS and seven single-slide clustering methods. We used the true cluster number as input for most clustering methods, except Seurat and Stardust because of their structure design.

### 3.1 Evaluation on simulation data

We performed comprehensive simulations to assess the performance of the 10 clustering methods (Fig 2). Details of the simulation procedure are provided in the “Technical details” section. In brief, we generated two datasets: one with and one without batch effects and one without batch effects, each comprising 4 slices with approximately 500 spots and 500 genes per slice (Fig 2a, d). The slices were divided into 4 layers, with varying mixture ratios of 3 cell types (mimicking eL2/3, eL4, eL5 in the STARmap mouse cortex data) to introduce cluster architecture within each slice. The UMAP visualizations of the gene expression data for both simulations highlight the differences between layers and slides (Fig 2b, e). For simulation 1 (Fig 2a-c), batch effects were introduced in the gene expression data. UMAP plots from simulation 1 and simulation 2 reveal distinct and expected patterns (Fig 2b, e): clear presence of batch effects in the 4 slices from simulation 1 only. Additionally, we introduced variation in the spatial coordinates of layer assignment between slices by applying a transformation to the spatial coordinates in slice 2, allowing us to evaluate the methods’ ability to handle potential shifts in spatial positioning. To comprehensively assess the clustering performance, we perform three sets of analyses: (1) single: single-slide clustering for each of the 4 slices separately, (2) multi1234: multi-slide clustering for all 4 slices simultaneously, and (3) multi134: multi-slide clustering for 3 slices excluding the slice with the coordinate transformation (slice 2). We note that in order to observe the impact of batch effects, we did not apply Harmony (for BASS, we removed the default step) in both simulations.

**Fig. 2.**
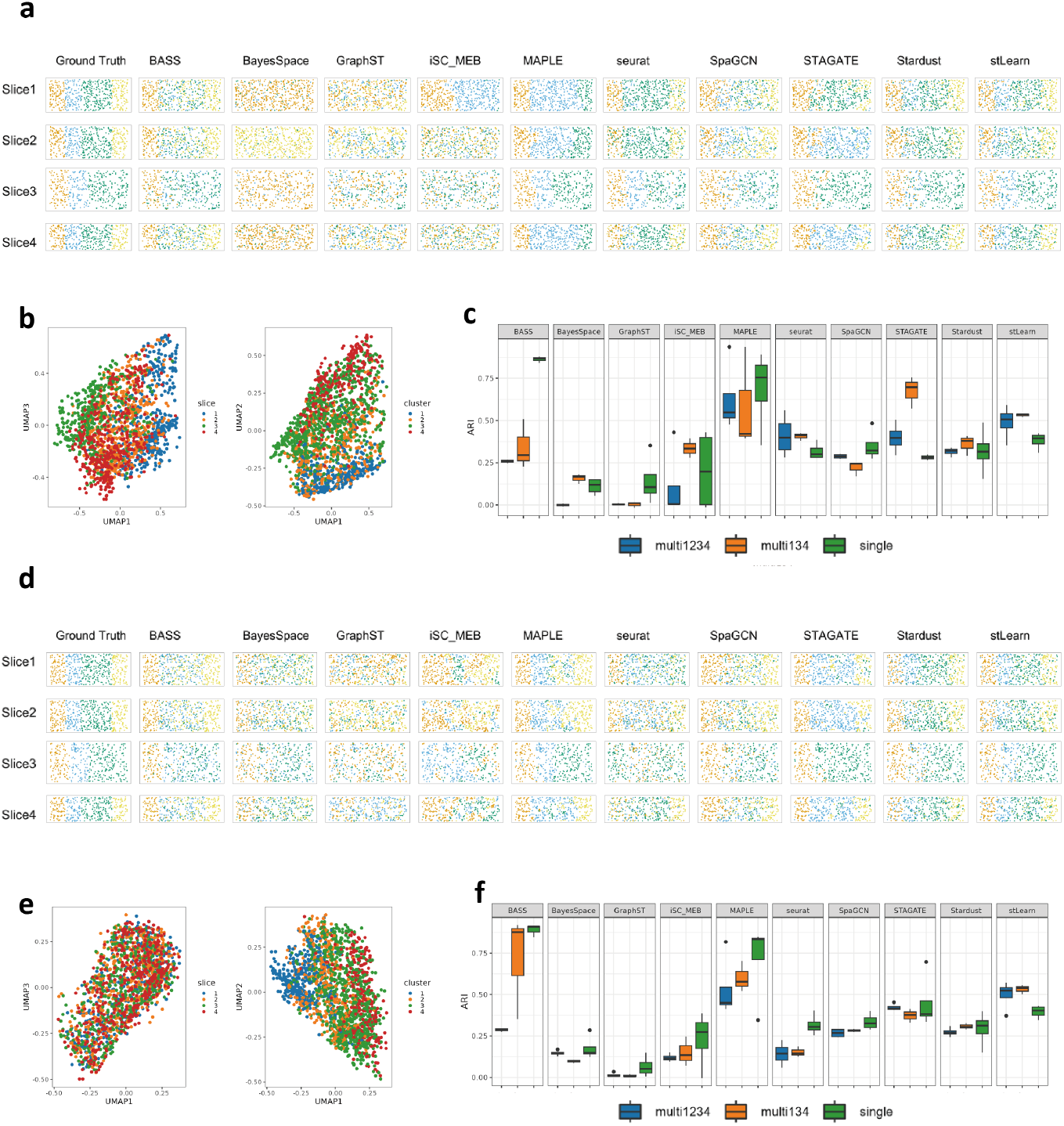
Evaluation on simulation data. With batch effect: **a**. Multi-slide clustering (multi1234) results and ground truth on simulation data1 slice1-slice4. **b**. Umap of simulation data1 cataloged with slice number and truth clusters. **c**. Boxplot of ARI for all the 10 methods on 4 sections on simulation data1. Without batch effect: **d**. Multi-slide clustering (multi1234) and ground truth on simulation data2 slice1-slice4. **e**. UMAP of simulation data2 cataloged with slice number and true cluster label. **f**. Boxplot of ARI for all the 10 methods on 4 sections on simulation data2.

For simulation 1 (Fig 2c), BASS with single-slide clustering performed the best (median ARI = 0.867), and MAPLE with single-slide clustering was the second best (median ARI = 0.754). Seurat, STAGATE, Stardust, and stLearn performed better in multi1234 and multi134 settings compared to single-slide clustering. BayesSpace and iSC.MEB showed improved performance in multi134 but decreased ARI in multi1234, suggesting their suitability for multi-slide clustering without coordinate variation. BASS, GraphST, MAPLE, and SpaGCN performed worse in both multi1234 and multi134 settings compared to single-slide clustering, indicating their sensitivity to batch effects. The performance of BayesSpace was also affected by batch effects, detecting each slide as a layer in multi1234 analysis. Most methods performed better without the coordinate transformation when multi-slide clustering, except MAPLE and SpaGCN. In the multi1234 analysis (Fig 2a), nearly all methods failed to distinguish layer 2 and layer 3, as well as layer 3 and layer 4. Specifically, BayesSpace and GraphST failed across almost all layers for every slice, while iSC.MEB only distinguished two layers in slice 1. Additionally, MAPLE, Seurat, and STAGATE tended to treat layer 2 and layer 3 as a single layer. In the multi134 analysis (Supplementary Figure 1c), BayesSpace and iSC.MEB were able to divide the samples into two layers across the 3 slices, STAGATE differentiated layer 2 and layer 3, while BASS only distinguished layer 1 from the other layers.

For simulation 2 (Fig 2f), BASS with single-slide clustering performed the best (median ARI= 0.910), and BASS with multi134 the second (median ARI = 0.877). stLearn performed better in multi1234 and multi134 compared to single-slide clustering. Most methods achieved their best performance in single-slide clustering and mostly performed better in multi134 than multi1234, suggesting their suitability for multi-slide clustering without coordinate variation. In the multi1234 analysis (Fig 2d), most methods still failed to distinguish layer 2 and layer 3, as well as layer 3 and layer 4. Again GraphST failed across all layers for every slice, iSC.MEB tended to treat layer 1 and layer 2 as a single layer, and STAGATE tended to merge layer 2 and layer 3. In the multi134 analysis (Supplementary Figure 2c), iSC.MEB was able to distinguish layer 1 and layer 2 across all 3 slices.

Comparing the ARI between simulation 1 and simulation 2 (Fig 2c, f), the performance of all methods was similar under single-slide clustering. BASS, BayesSpace, STAGATE, iSC.MEB, and stLearn performed better in simulation 2 than simulation 1 in the multi1234 setting, suggesting improved performance without batch effects. BASS, MAPLE, SpaGCN, and stLearn exhibited better performance in simulation 2 than simulation 1 in the multi134 setting, indicating their improved efficiency without batch effects and coordinate variation. Examining specific clusters in the presence of coordinate variation (Supplementary Figure 2b), BayesSpace could distinguish layer 1 from other layers without batch effects but failed to separate any layers with batch effects. SpaGCN encountered greater difficulty in differentiating layer 3 and layer 4 even without batch effects. In the absence of coordinate variation (Supplementary Figure 2c), BASS demonstrated significant improvement, suggesting its superior performance without batch effects and coordinate variation.

### 3.2 Evaluation on the human dorsolateral prefrontal cortex (DLPFC)

Moving to real data, we evaluated the clustering methods using the human DLPFC data from the 10x Visium platform, comprising 12 slices from t 3 independent adults [15] (Fig 3). The samples were sequenced with a median depth of 291.1 *×* 106 reads, resulting in an average of 3,462 spots per slide and 1,734 genes per spot. The data was annotated by pathologists into 7 layers (L1-L6 and white matter (WM)) (Fig 3a). The 12 samples were divided into 3 groups (group 1: 151507-151510, group 2: 151669-151672, group 3: 151673-151676) based on morphology similarity. We performed single-slide clustering individually on the 12 slides and conducted multi-slide clustering within the 3 groups using both single-slide and multi-slide methods. For a fair comparison in the evaluation for multi-slide clustering with single-slide clustering methods, in addition to simply pooling all the samples together, we also evaluated their performance with the integration of PASTE [37] for spatial coordinate alignment and Harmony [38] for batch effect removal as a pre-processing step.

**Fig. 3.**
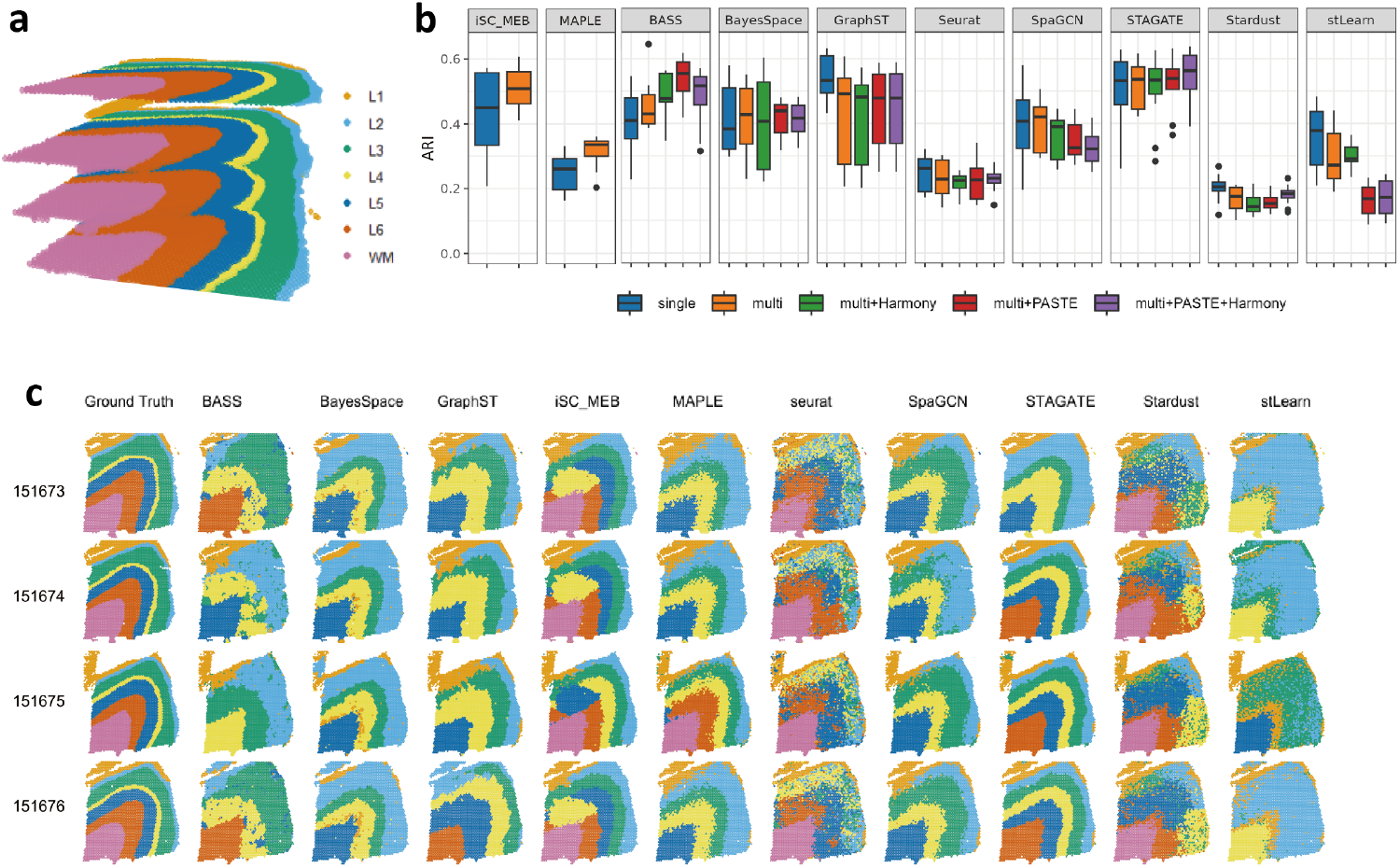
Evaluation on DLPFC. **a**. Ground truth of tissue regions for slides in group 3 (samples 151673- 151676). Spatial coordinates are aligned using PASTE. **b**. Boxplot of ARI for all 10 methods on 12 sections. For each method, we conduct both single-slide and multi-slide clustering. With or without PASTE or Harmony for multi-slide clustering, we consider four procedures: multi (neither PASTE nor Harmony), multi+Harmony, multi+PASTE, multi+PASTE+Harmony. **c**. Clustering results from all 10 methods in the multi-slide clustering case on group 3 (multi, samples 151673-151676).

We start with a comprehensive evaluation of all methods applied to single-slide clustering. STAGATE demonstrated the greatest ARI (median = 0.558), followed by GraphST with the second highest ARI (median = 0.533) (Fig 3b), indicating the superior performance of the graph network structure. The Bayesian models, BASS (ARI: median = 0.437), BayesSpace (ARI: median = 0.382), and iSC.MEB (ARI: median = 0.450) also performed well, with difference within reasonable bounds observed across the three methods. For multi-slide clustering, most non-Bayesian single-slide methods exhibit inferior performance compared to their single-slide clustering ARI. Bayesian modeling methods, BASS (median ARI = 0.449), BayesSpace (median ARI = 0.428), and iSC.MEB (median ARI = 0.508) typically performed better in multi-slide clustering scenarios. BayesSpace provided a clearer distinction between L3 and L5 within group 3 than other methods (Fig 3e, Supplementary Figure 4). The inferred boundaries became hazy when GraphST was used for multi-clustering. STAGATE’s capacity to segment L3, L4, and L5 decreased when applied to group 3 (especially slide 151673, Supplementary Figure 4). STAGATE (median ARI = 0.534) had the best ARI, whereas iSC.MEB (median ARI = 0.508) had the second highest ARI. Maple had better performance in multi-slide clustering (median ARI = 0.336) compared to single-slide clustering (median ARI = 0.226), although the ARI remained in the relatively moderate range. Comparatively, Seurat and Stardust provided slightly noisier layer predictions.

**Fig. 4.**
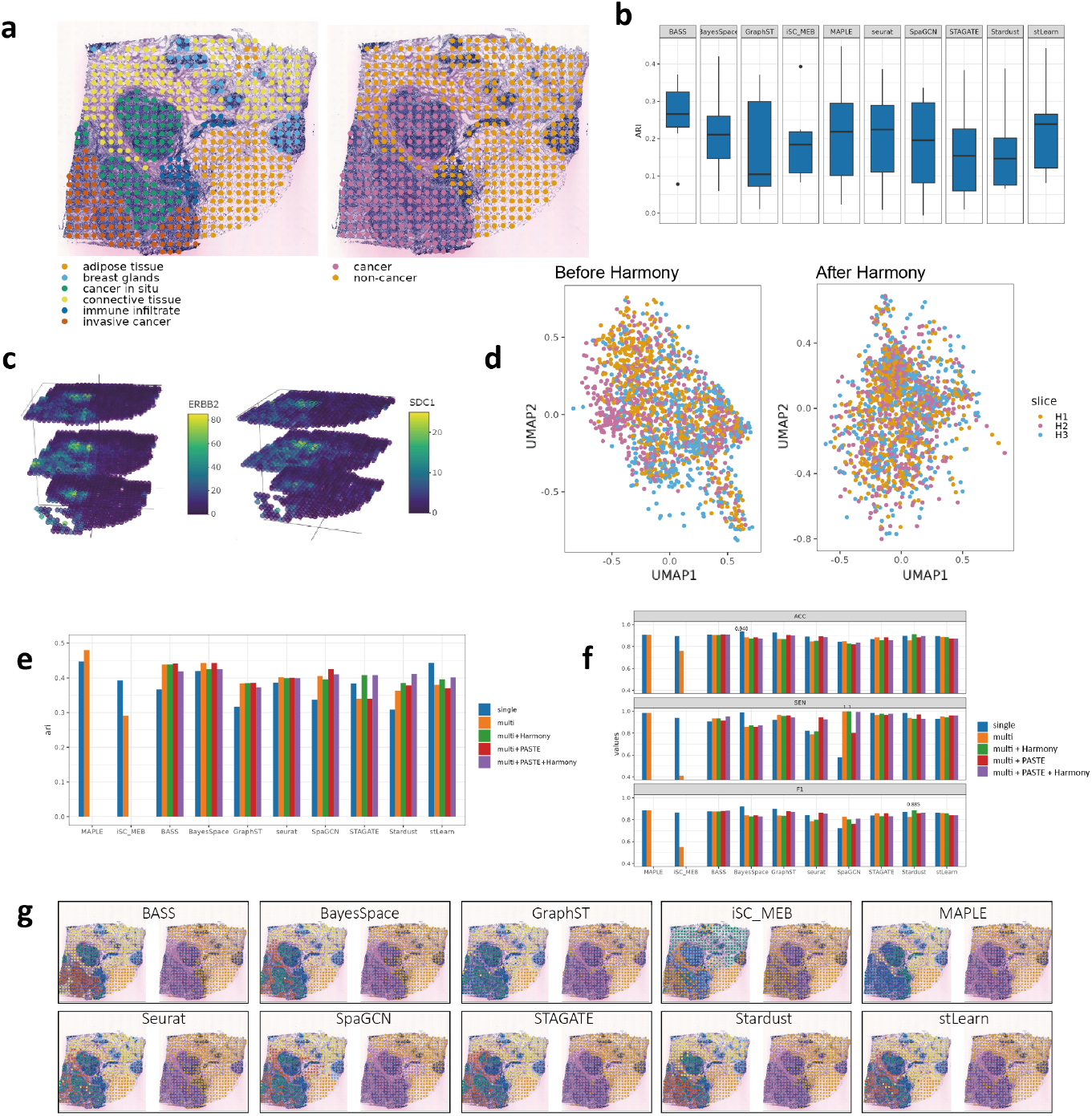
Evaluation on HER2+ breast cancer data. **a**. Left: Ground truth of tissue regions on H1. Right: cancer regions (including both cancer in situ and invasive cancer regions) and non-cancer region on H1. **b**. Boxplot of ARI for all the 10 methods on 8 sections (A1-H1). We used each method to perform single-slide clustering. **c**. gene expression of ERBB2 and SDC1 in H1-H3. **d**. UMAP of gene expression before and after Harmony (H1-H3). **e**. Line chart of ARI on H1. We conducted analyses in five manners: single, multi(neither Harmony nor PASTE), multi+Harmony, multi+PASTE, multi+PASTE+Harmony. Note that neither was added to MAPLE and iSC.MEB. **f**. Dodge bar plot for accuracy, sensitivity, and F1-score to evaluate the ability to detect cancer regions in 5 cases. **g**. H1 clustering results in the multi-slide clustering case.

Next we evaluated how the pre-processing procedures impacted clustering performance in multi-slide clustering scenarios for single-slide clustering methods. Harmony had little effect on (Fig 3b), the performance of GraphST (median ARI = 0.491 vs 0.491 with and without Harmony), Seurat (median ARI = 0.221 vs 0.227 with and without Harmony), Stardust (median ARI = 0.175 vs 0.175 with and without Harmony) and STAGATE (median ARI = 0.537 vs 0.537 with and without Harmony), and impaired the performance of BayesSpace (median ARI = 0.407 vs 0.473 with and without Harmony),and SpaGCN (median ARI = 0.390 vs 0.420 with and without Harmony. However, when Harmony is applied to BayesSpace, it can provide greater clarity about the distinction between L5 and L6 in group 3 (Supplementary Figure 5). Interestingly, PASTE negatively affected the performance of most methods, except STAGATE (median ARI = 0.561 vs 0.537 with and without PASTE). With both PASTE and Harmony, only Seurat (median ARI = 0.229 and 0.228 with and without PASTE and Harmony), Stardust (median ARI = 0.215 and 0.175 with and without PASTE and Harmony), and STAGATE (median ARI = 0.566 and 0.537 with and without PASTE and Harmony) showed improved ARI. Although ARI failed to improve for most methods, all methods yielded better intra-group similarity with the application of PASTE and Harmony (Supplementary Figure 7). STAGATE consistently outperformed other methods, particularly when combined with PASTE and Harmony in multi-slide scenarios (median ARI = 0.566 vs ¡=0.510 for other methods). In addition, since Harmony is part of BASS’s default procedure, we also tried applying both PASTE and Harmony with BASS. BASS with Harmony (ARI: median = 0.477) shows an increase compared to BASS without preprocessing techniques (ARI: median =0.449), and BASS with PASTE (ARI: median = 0.555) has the highest performance through all the scenarios of BASS.

**Fig. 5.**
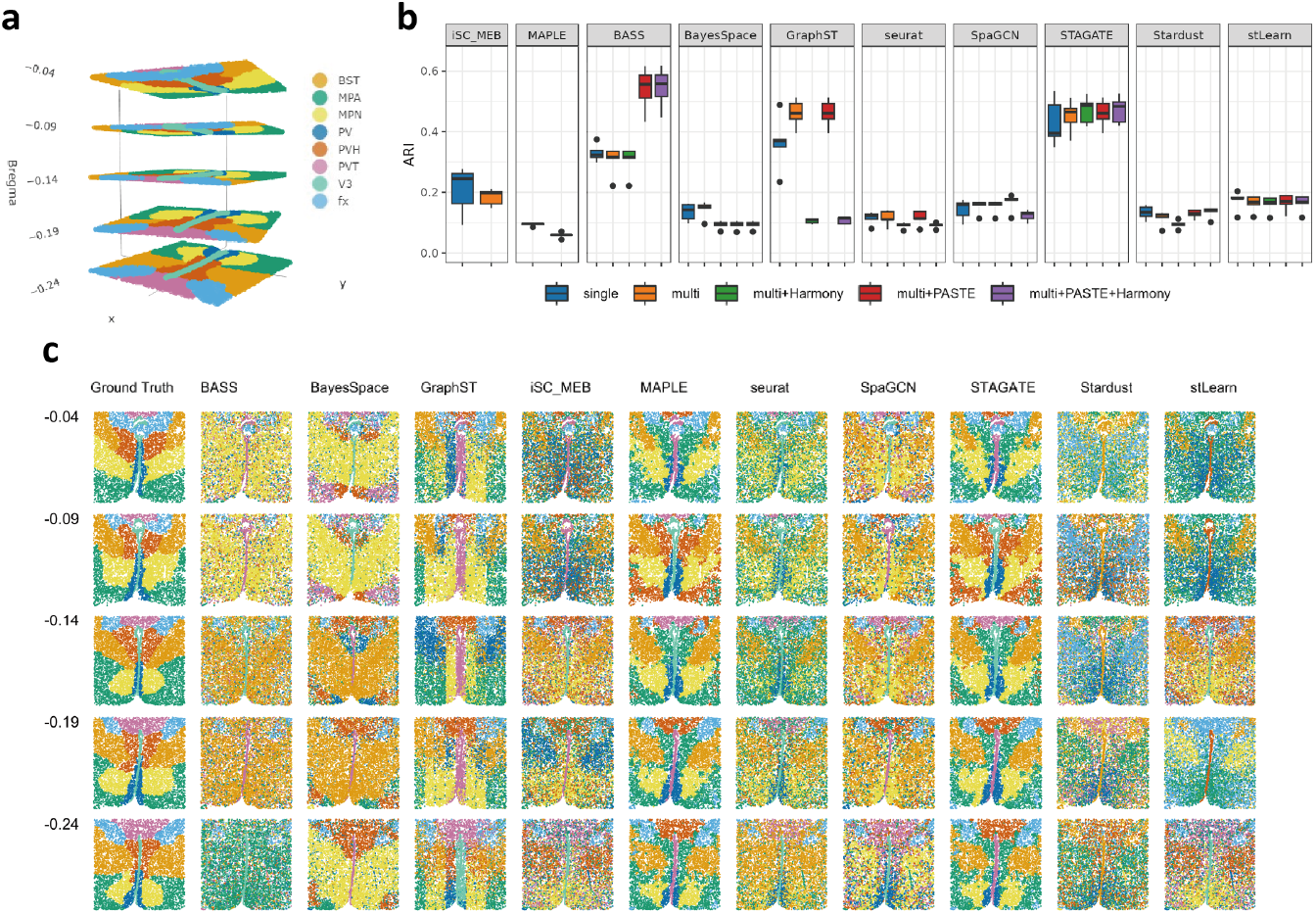
Evaluation on mouse hypothalamus MERFISH data. **a**. Ground truth of tissue regions applied with PASTE on mouse hypothalamus MERFISH data(Bregma -0.04,-0.09,-0.14,-0.19,-0.24). **b**. Boxplot of ARI for all the 10 methods on 5 sections. For each method, we performed both single-slide and multi-slide clustering. For multi-slide clustering, we conducted analyses in 4 manners: multi(neither PASTE nor Harmony), multi+Harmony, multi+PASTE, multi+PASTE+Harmony on BASS and all seven single-slide clustering methods. **c**. Multi-slide clustering results.

### 3.3 Evaluation on HER2+ breast cancer data

To further evaluate clustering performance, we analyze the efficacy of clustering methods on the HER2+ breast cancer data (Fig 4). ST data on HER2+ tumors was collected from eight individuals (patients A-H). We choose one slice per patient with pathologist annotation for analysis (A1, B1, C1, D1, E1, F1, G2, H1). We choose three slices (H1-H3) from patient H for multi-slide clustering. PASTE and Harmony are utilized for single-slide methods in the context of multi-slide clustering.

For single-slide clustering (Fig 4b), we observe that BASS performs the best (median ARI = 0.266), followed by stLearn (median ARI = 0.239). Some models with complex structures show relatively worse performance in this data (median ARI = 0.104 and 0.154 for GraphST and STAGATE). In A1, virtually all sections in the middle are invasive cancer regions, but BayesSpace and iSC.MEB separate them into two regions, which could be due to the similarity in invasive cancer and connective tissue (Supplementary Figure 8). It’s challenging for all the approaches to separate cancer in situ, breast glands, and adipose tissue in B1, C1, D1, E1, and F1. Most methods can delineate the adipose cluster in G2 but fail with other clusters.

For multi-slide clustering (Fig 4c), we only evaluate the ARI in H1 since there is no ground truth available for H2 and H3. MAPLE (ARI = 0.480) is the best performer, followed by BASS (ARI = 0.439). Most methods have better performance in multi-slide clustering compared to single-slide clustering in H1, with the exception of STAGATE, iSC.MEB and stLearn. Most single-slide clustering methods perform at least the same or better with PASTE. With the incorporation of Harmony, STAGATE, Stardust, and stLearn perform better. GraphST and Seurat are minimally impacted by these two preprocessing procedures. SpaGCN performs best with PASTE (ARI = 0.425), while Stardust performs best with both PASTE and Harmony (ARI = 0.412). For most methods, multi-slide clustering with PASTE only is recommended. It remains difficult for all the clustering methods to separate cancer in situ and invasive cancer (Supplementary Figure 9), which are similar in the expression of cancer marker genes (GLYATL2, FNBP1L, SDC1). The division of cancer areas by BayesSpace, Seurat, SpaGCN, and STAGATE closely resembles ground truth. However, detecting the connective tissue between the two cancer areas in H1 is extremely difficult.

A major question that we are concerned about is whether a clustering method can separate cancer from non-cancer areas. We annotated the clustering result based on cell-type marker genes to compare the identification capabilities of different methods in H1-H3 (Fig 4e, Methods). Clusters with high mean expression of cancer marker genes GLYATL2, FNBP1L, and SDC1 are labeled as cancer regions. To evaluate the performance, we defined ground truth based on pathologist’s annotations, with the pathologist annotated cancer in situ and invasive cancer region as cancer regions, while others as non-cancer region (Fig 4a). We computed accuracy (ACC), sensitivity, and F1-score to evaluate the performance of various methods (Fig 4d). This comparison also involves the application of single-slide methods for multi-slide clustering. Stardust with PASTE consistently demonstrates the highest accuracy (ACC=0.915) and F1-score (F1=0.885) in multi-slide clustering. The sensitivity of SpaGCN reaches 1 in the case without preprocessing technique and the case with Harmony, demonstrating precise distinction of color where cancer regions appears to be darker and denser in the H&E staining histology image. In H2 and H3, iSC.MEB, Seurat, SpaGCN, STAGATE, and stLearn provide a more precise delineation of the cancer regions (Supplementary Figure 9-10). In H1 and H2, methods like MAPLE, SpaGCN, STAGATE, and stLearn tend to classify the immune infiltrate as the cancer area, perhaps because of its dark color.

### 3.4 Evaluation on Mouse hypothalamus MERFISH data

Finally, we evaluated the 10 methods on mouse hypothalamus MERFISH data. The data was obtained by dissecting a rostral portion of the mouse hypothalamus containing the preoptic region and surrounding nuclei (*∼*2.5 mm *×* 2.5 mm *×* 1.1 mm, Bregma +0.5 to *−*0.6). We selected 5 adjacent tissue sections: Bregma -0.04, -0.09, -0.14, -0.19, and -0.24 mm from a consecutive brain hypothalamic region of one animal, which is a core area controlling many social behaviors and homeostatic functions. We utilized the manually annotated spatial domains from the BASS paper as the ground truth, which included the third ventricle (V3), bed nuclei of the stria terminalis (BST), columns of the fornix (FX), medial preoptic area (MPA), medial preoptic nucleus (MPN), periventricular hypothalamic nucleus (PV), paraventricular hypothalamic nucleus (PVH), and paraventricular nucleus of the thalamus (PVT) (Fig 5a). For each method, we performed both single-slide and multi-slide clustering and further compared the performance of Harmony and PASTE combined with the BASS method and all seven single-slide clustering methods.

Comparing the ARI of single-slide and multi-slide clustering without data preprocessing (Fig 5b), we found that the highest ARI was achieved by STAGATE under multi-slide clustering (median ARI = 0.466), and the second highest was GraphST, also under multi-slide clustering (median ARI = 0.462). Besides STAGATE and GraphST, BayesSpace and SpaGCN also performed better with multi-slide than with single-slide clustering. However, some methods designed for multi-slide clustering did not improve with multi-slide clustering, possibly due to batch effects. For example, BASS (median ARI = 0.324 and 0.317 single and multi-slide clustering). In terms of discriminating power (Fig 5c, Supplementary Figure 11a, b), most methods successfully distinguished V3 and fx. iSC.MEB delineated BST in single-slide clustering but merged it with PVH and MPN in multi-slide clustering. BASS tended to divide MPN and MPA in a different pattern, which actually is consistent with the MPA and LPO clusters labeled in the original paper for this MERFISH data (Fig 5B in [36]). GraphST could even distinguish the cluster ACA into several part only labeled in the original paper [36]. GraphST and STAGATE performed well but tended to merge MPA and PVH in multi-slide clustering.

Assessing the impact of PASTE and Harmony on multi-slide clustering, we found that BASS (median ARI = 0.317) improved significantly with PASTE (median ARI = 0.577) and performed best with both PASTE and Harmony (median ARI = 0.578). In contrast, BayesSpace(median ARI = 0.097 vs 0.151 with and without Hamorny and PASTE) performed worse with either added. GraphST (median ARI = 0.110 vs 0.462 with and without Hamorny) and, Seurat (median ARI = 0.111 vs 0.114 with and without Hamorny) improved slightly with PASTE but performed much worse with Harmony. SpaGCN (median ARI = 0.176 vs 0.163 with and without PASTE) improved substantially with PASTE. STAGATE did not change significantly with the two techniques. In terms of discriminating power (Supplementary Fig 12a, b, c), BASS with PASTE could now distinguish MPN and MPA. GraphST performed significantly worse with Harmony. Most methods exhibited little change after using PASTE or Harmony.

## 4 Discussion

As a pivotal part of ST data analysis, clustering facilitates the exploration of intercellular interactions across diverse spatial locations and the identification of genes exhibiting spatial expression patterns that transcend individual cells. Although most ST clustering algorithms are trained independently on individual tissue sections, the advent of multi-slide clustering methods has emerged due to the availability of numerous contiguous tissue sections derived from the same or similar tissue specimens within or across individuals, sharing comparable spatial domains and cell type compositions. Leveraging these conditions can potentially enhance the performance of clustering across different samples. This forms the central focus of investigation in the paper.

This study benchmarks the performance of 10 clustering methods for multi-slide ST data using two simulated datasets and three real datasets. BASS performed best in single-slide clustering, with significant improvements when batch effects were removed and coordinate transformation was applied. In multi-slide clustering, most methods, except SpaGCN, showed better performance without coordinate transformation. For the DLPFC data, STAGATE outperformed other methods, while Bayesian models also demonstrated good performance, with multi-slide clustering generally enhancing results. However, discerning cortex layers from histological images remained challenging. In HER2+ breast cancer data, BASS excelled in single-slide clustering, while stLearn performed well in multi-slide clustering, especially when incorporating image data. Although methods like BayesSpace and iSC.MEB performed well when spot locations were arranged in an equidistant lattice, they struggled with more irregular arrangements. In contrast, STAGATE and GraphST handled non-uniform spot locations effectively in the mouse hypothalamus data, suggesting that graph-based models are more robust to coordinate transformations. While specifying the number of clusters a priori improves performance for most methods, exceptions like Seurat and Stardust do not require this step, emphasizing the need for flexibility in determining cluster numbers for novel datasets.

We evaluated the impact of two data preprocessing techniques, Harmony and PASTE. PASTE worked well when coordinate transformation was substantial, while Harmony’s balancing approach sometimes conflicted with model assumptions or image-derived features, as seen in methods like BayesSpace and SpaGCN. Specifically, Harmony had minimal impact on most methods, benefiting BASS but reducing BayesSpace’s performance. In contrast, PASTE significantly improved BASS, particularly in the presence of coordinate transformation, while negatively affecting SpaGCN on the DLPFC dataset and enhancing its performance on breast cancer and mouse hypothalamus data. When applying both Harmony and PASTE, some methods performed worse than without both techniques, potentially due to duplicate data balancing. These mixed outcomes suggest that the cooperation between preprocessing techniques and the underlying data structures is critical. Selecting an appropriate preprocessing technique depends on the method’s sensitivity to spatial transformations and data balancing.

It is important to note that ARI, although a best and convenient statistic to quantify the performance of clustering methods, does not contain all useful information. It should be combined with visual evaluation of clustering results for more comprehensive performance evaluation. However, clustering results that differ from existing ground truth does not necessarily indicate method inadequacy. In the real data of clustering, there may exist more than one valid solution, each representing a different shared characteristic, for example, a different mixture of cell types. The unsupervised clustering could yield other results which may also be reasonable.

Potential factors influencing the performance of multi-slide clustering include inherent structure design of the clustering method itself, batch effects, coordinate transformations, ST technologies employed for data acquisition, and the dimension of the overall feature matrix. The first three factors have been elucidated in the preceding paragraph. Occasionally, abnormalities can arise due to the ST technologies employed for data acquisition not compatible with the methods designed, resulting in suboptimal performance. For example, methods such as BayesSpace and iSC.MEB encounter difficulties in identifying neighboring spots when spot locations deviate from a non-equidistant lattice arrangement. Although iSC.MEB provides alternative techniques for finding neighbors beyond the 10X platform, these techniques can be time-consuming and may still fail to identify neighbors for most spots. Furthermore, iSC.MEB can sometimes encounter issues with loop invocations and cause error when running, which can be mitigated by reducing the maximum item count, albeit at the cost of reduced accuracy in the final results. Concerning the final factor, feature matrix dimension, consider stLearn as an example, which utilizes ResNet50 for feature selection and has a limited capacity to process the input matrix. stLearn exhibits enhanced performance when the total number of spots and genes are smaller, thus, making it more suitable for single-slide scenarios than multi-slide scenarios with highdimensional feature matrix. stLearn is not the only method that has dimensionality constraints: most methods have their respective capacity ranges, which should be taken into consideration when selecting the appropriate clustering method for a specific dataset. The techniques employed for dataset alignment or harmonization are not universally suitable for all datasets or all clustering methods. Although more evaluations are still warranted when encountering new situations, our study provides useful guidelines for multi-slide clustering of ST data from the most comprehensive evaluations to date.

## 5 Technical details

### 5.1 Adjusted Rand index (ARI)

ARI is a metric used to quantify the similarity between two clusterings. In our study, we utilized the ARI function from the ‘aricode’ R package to calculate the ARI values. ARI has an expected value of 0 in the case of a random partition and takes a value of 1 in the case of perfect agreement. Given a set S of n elements and two groupings (e.g., results from a clustering method and ground truth) of these elements, denoted as *X* = *X*_1_, *X*_2_, …, *X*_*r*_ and *Y* = *Y*_1_, *Y*_2_, …, *Y*_*s*_, respectively, the agreement between X and Y can be summarized in a contingency table [*n*_*ij*_], where *n*_*ij*_ represents the number of elements that belong to both cluster *X*_*i*_ and cluster *Y*_*j*_.

Then the Adjusted Rand Index is,

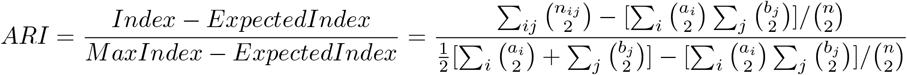

### 5.2 PASTE

PASTE is a method that makes probabilistic alignment of ST data. It aligns and integrates ST data from multiple adjacent tissue slices. For input of PASTE, we pooled the slices of ST data into different groups according to their spatial locations. After that, we sort the matrix with ids expressed in layer data. Meanwhile, we replace ids with their corresponding spatial coordinates. Finally, we ran PASTE to align slices and generated a 3D plot in R to see what the result was.

### 5.3 Harmony

Harmony is a widely-used batch effect correction method. We used Harmony (1.2.0) in our analysis. Following the required steps before running Harmony, we first obtained log normalized counts using logNormCounts() on single-cell experiment objects. Next, we ran PCA using runPCA(), and finally used RunHarmony to perform batch effect correction across different slices.

### 5.4 Simulation Design

We simulated two datasets: one with batch effects and one without. Both datasets comprised 4 slices, each containing 500 genes and approximately 500 spots. Each slice consisted of 4 layers with varying mixture ratios of cell types. The spot positions were simulated randomly with a uniform distribution within a specified area (250*100 pixels, layer1: x in [0,50); layer2: x in [50,100); layer3: x in [100,200); layer4: x in [200,250)). For slice 2, we added 50 pixels to both the x and y coordinates to mimic parallel slice movements. To further introduce variations, we removed 20% of the total area on the right for slice 3 and 20% of the total area at the bottom for slice 4.

Regarding gene expression, we first estimated parameters for 3 cell types (eL2/3, eL4, eL5) from a real STARmap mouse cortex data using the R package splatter. We then simulated several single-cell gene expression count matrices using these cell types with or without batch effects for 4 slices. Subsequently, we assigned cells to different spots, with each spot containing 10 cells. The primary variation across different layers was arise from the fluctuation of the ratio of eL2/3 and eL4, with eL5 representing a random effect. Our layers were arranged horizontally with varying mixture ratios of the 3 cell types: layer 1 (eL2/3:0.6, eL4:0.3, eL5:0.1; x-pixel scale: 0-50), layer 2 (eL2/3:0.5, eL4:0.4, eL5:0.1; x-pixel scale: 50-100), layer 3 (eL2/3:0.4, eL4:0.5, eL5:0.1; x-pixel scale: 100-200), and layer 4 (eL2/3:0.3, eL4:0.6, eL5:0.1; x-pixel scale: 200-250).

### 5.5 Refine clusters

This method, developed specifically for clustering analysis, has been detailed by Chen et al. [39] in the supplementary notes.

### 5.6 Annotation for clusters with cell-type marker gene

If the ground truth is not provided, we can perform annotation with cell-type marker genes, for the primary cell type within each cluster. In detail, suppose cell type a is the most abundant cell type in cluster A, we calculate the mean gene expression of marker genes for cell type a for all the existing clusters, and annotate the cluster with the largest mean gene expression as cluster A. When two clusters share the same predominant cell type, we compared the mean expression of marker genes in the two clusters and created a plot withspatial coordinates to assign the annotation manually. If there is a region with one refined cluster, but two original clusters from results, it is because the clustering method originally divided more than supposed categories when clustering, and one of them was annotated to cancer the other didn’t. The refined cluster separates the two clusters into one. When evaluating on HER2+ breast cancer data, GLYATL2, FNBP1L, SDC1 are the cancer marker gene [35]. We employ a threshold for the mean gene expression at each spot to distinguish between cancer and non-cancer regions. We use 3 as the threshold through manual observation through H1-H3. Regions with gene expression levels exceeding this threshold are classified as cancer regions. Annotation using cell-type marker genes could be vulnerable to misclassification, thus we also offer the raw cluster data for reference.

### 5.7 Other technical details

Seurat adheres to a set of fundamental steps. The SCTransform function is first utilized to standardize the data and identify significant variables, in which a subsample of 5000 cells is employed to construct the negative binomial regression model. The features will be found by the residual variance from the negative binomial regression model. Then Principal Component Analysis (PCA) is performed to reduce the dimensionality, resulting in top 50 principal components (PCs). Subsequently, the initial 30 principal components are used to identify neighboring data points in the K-Nearest Neighbors (KNN) algorithm. Finally, the Louvain approach is adopted to identify the clusters.

For stLearn, we follow the stSME clustering tutorial including a novel normalization method. We apply a filter to select genes with non-zero counts in ¿=2 cells, and perform Umi normalization and log1p transformation. We segment H&E images into small tiles according to the spatial location of spots and extract the underlying morphological traits. To normalize the data, we utilize spatial location (S), tissue morphological characteristic (M), and gene expression (E) information. We performed PCAto obtain the top 50 PCs. Finally, we apply the k-means algorithm to perform clustering.

For iSC.MEB, we use Seurat to read and preprocess input data. We employ Log-Normalization with a cell-level scale of 10000 and utilize local polynomial regression to identify 2000 features. We merge Seurat objects to form an iSC.MEB object, which involves utilizing ’SPARK-X’ to identify the top 2000 genes that exhibit spatial variability. In the process of generating neighboring data, we set ‘Visium’ for DPFLC and ‘ST’ for other datasets. When performing clustering, we select the top 15 PCs and designate the true cluster number as the cluster number. Typically, 25 iterations are executed, but occasionally there may be issues with loop invocations. In this scenario, we can only decrease the number of iterations.

We followed the vignettes of BASS v1.1.0.016. We used all default settings except the burnin, which we increased to 10000 iterations to ensure the convergence of spatial parameters. Also, there is a built-in option in BASS to perform batch correction through HARMONY, we turn on or off this option as described in the Results section. In the default setting of BASS, we used top 20 PCs with no other feature selection methods. Notably, BASS authors suggested using SPARK-X to extract top spatially variable genes in the DLPFC dataset, but we skipped this step for fairer comparison with other methods.

For BayesSpace, we followed its vignettes and used all default settings but increased the number of PCs from the default 15 to 50. We used the default number of 50000 iterations for MCMC. Note that BayesSpace only supports specific spatial distribution of spots in neighborhood detection. In the analysis for simulated data and mouse hypothalamus data by MERFISH, it failed to detect neighborhoods.

In our test for SpaGCN, we keep parameters the same as in the step-by-step tutorial given in GitHub repository except for max epochs. We change this parameter to a large number because we have sufficient computational power, and we suppose it will do no harm to the result in this unsupervised learning. The number of clusters is set based on the ground truth (manual annotation). When implementing the multi-slide tests, we slightly modify SpaGCN’s “calculate adj matrix” to allow the processing of multiple staining images since the original function can only input one image at a time. Specifically, the modified function processes RGB values from different images separately at first. These RGB values alongside spatial coordinates will then be used to compute pairwise distances between all spots in all slides.

## Author Contribution

Caiwei Xiong implemented four clustering methods, performed data preprocessing, simulation, analysis, visualization, and contributed to writing. Shuai Huang implemented three clustering methods. Muqing Zhou implemented one clustering method and maintained the GitHub repository. Yiyan Zhang implemented two clustering methods and performed the Harmony processing. Wenrong Wu performed the PASTE processing. Dr. Yun Li and Jiawen Chen provided guidance on method selection and supervision.

## Acknowledgments

We thank Li lab members for providing advice on method selection and feedback on the manuscript. Figure 1 is created via bioRender.

## Code avialability

All code used in this study can be found at https://github.com/zmqsonata/ST-multi-slide-clustering.

## Notes

### Competing Interest Statement

The authors have declared no competing interest.

