## Supplementary Figures for "A comprehensive comparison on clustering methods for multi-slide spatially resolved transcriptomics data analysis"

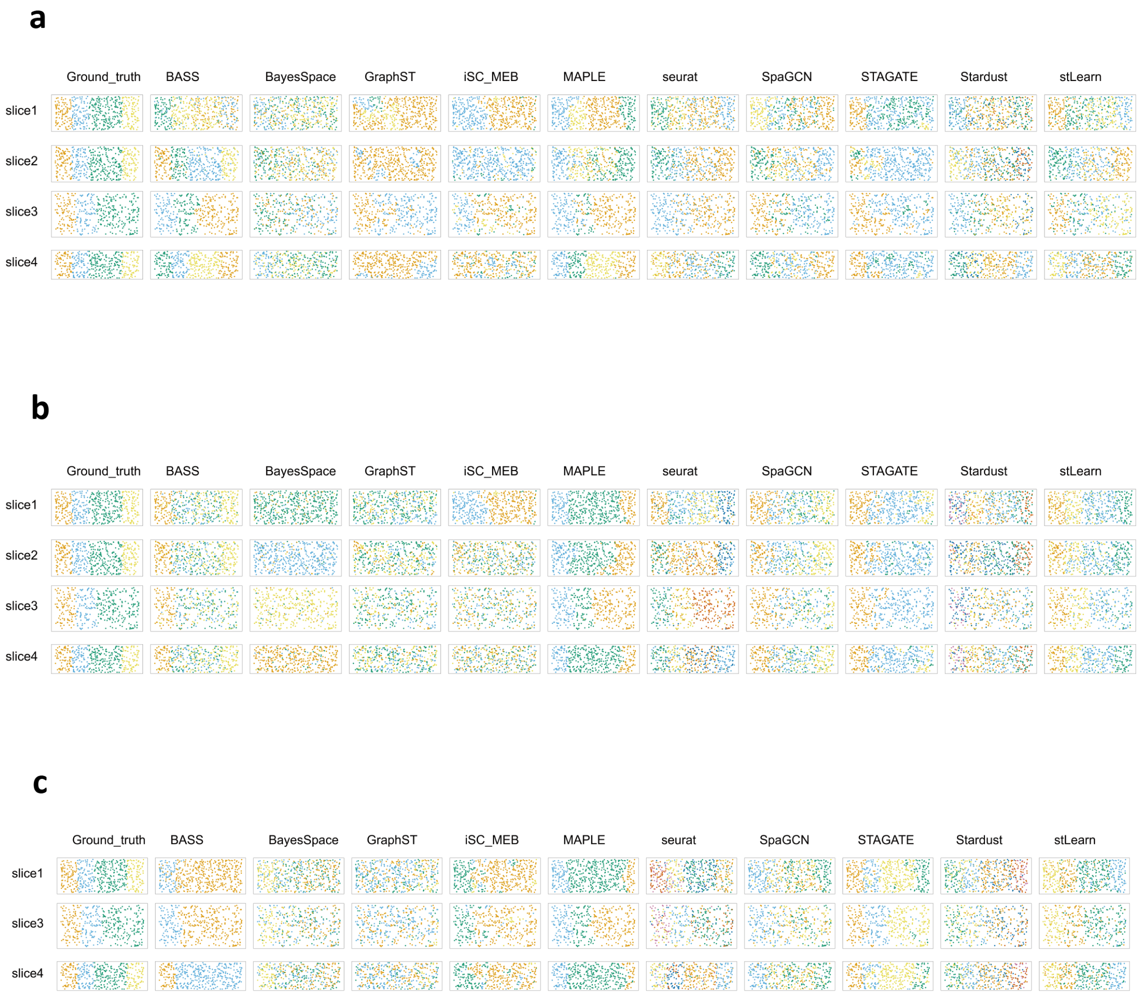


**Supplementary Figure 1.** 10 Clustering methods on simulation data 1 with batch effect. **a.** Single-slide clustering on slice1-slice4 separately. **b.** Multi-slide clustering on slice1-slice4 simultaneously. **c.** Multi-slide clustering on slice1-slice3 simultaneously.

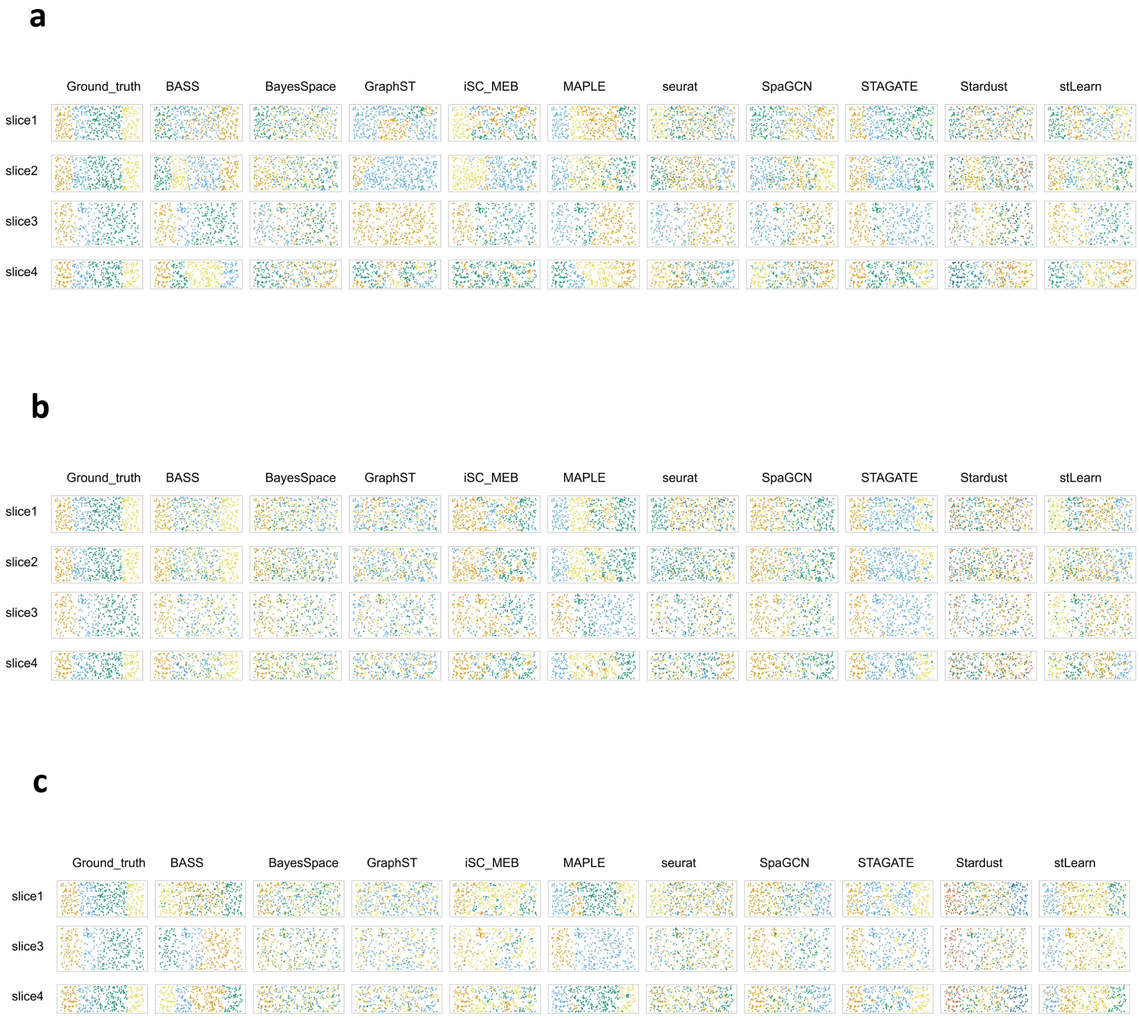


**Supplementary Figure 2.** 10 Clustering methods on simulation data 2 without batch effect. **a.** Single-slide clustering on slice1-slice4 separately. **b.** Multi-slide clustering on slice1-slice4 simultaneously. **c.** Multi-slide clustering on slice1-slice3 simultaneously.

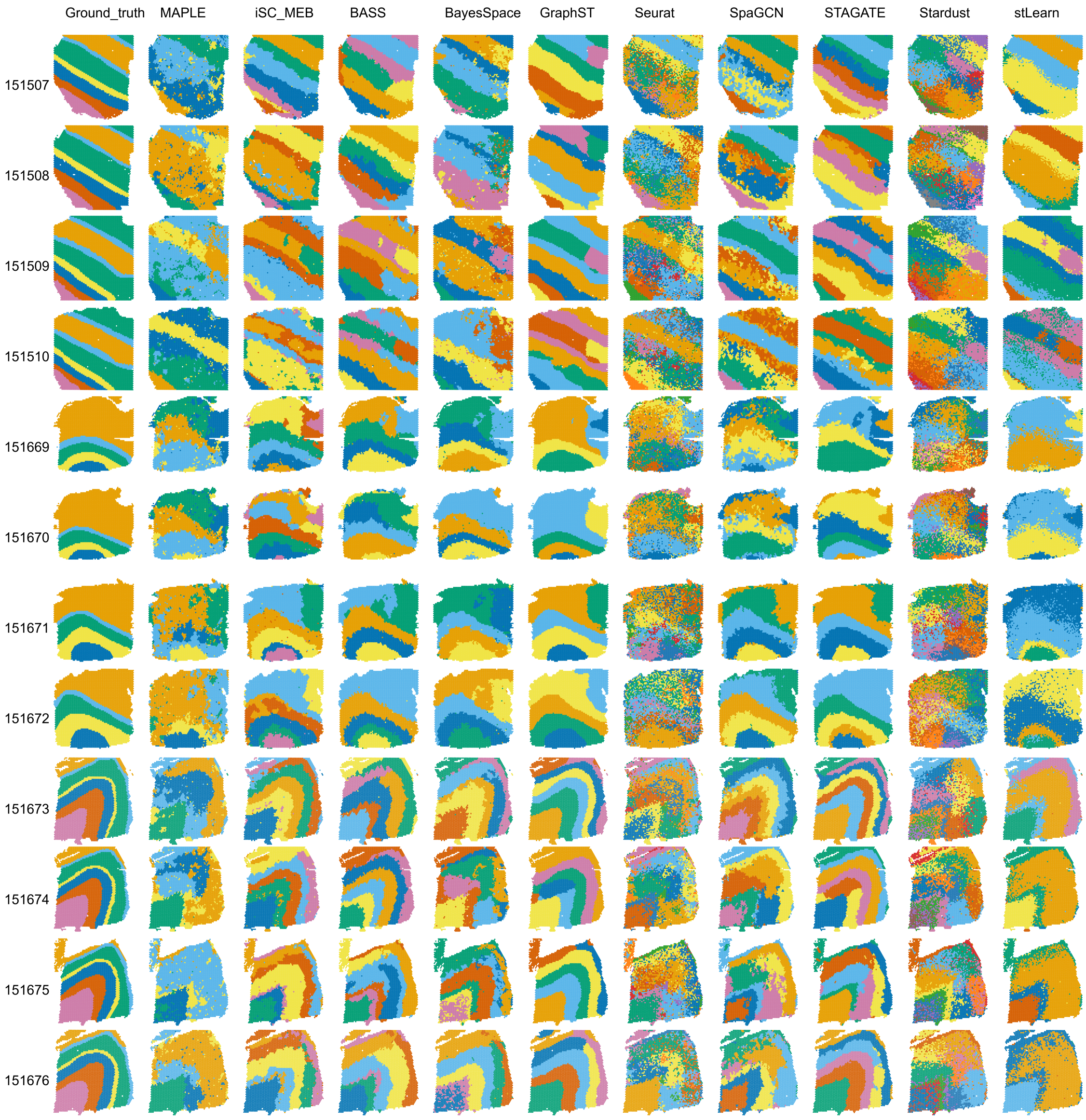


**Supplementary Figure 3.** Single-slide clustering results on DLPFC. Clustering of tissue locations based on all 10 methods in the single-slide clustering case across all 12 samples.

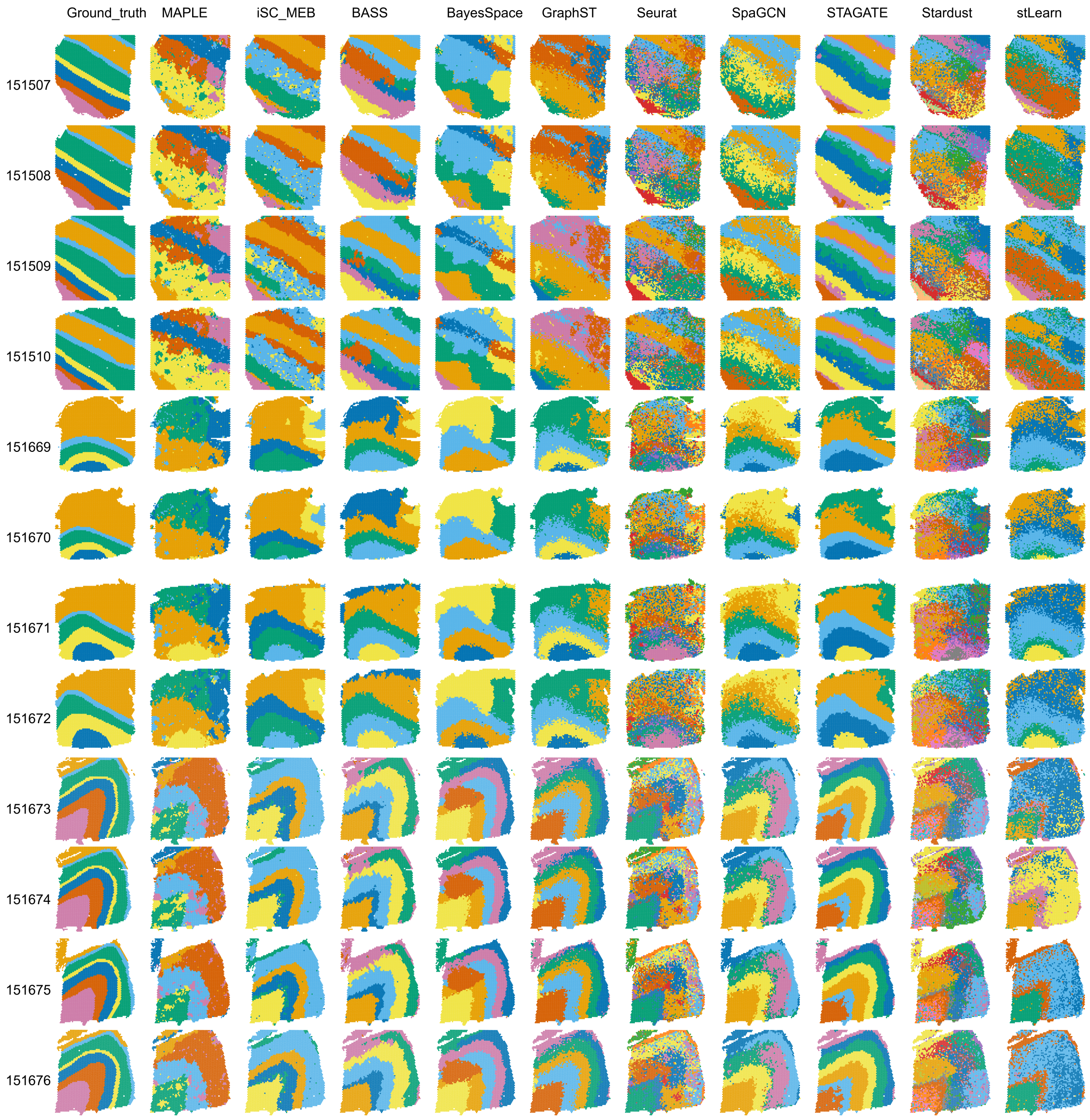


**Supplementary Figure 4.** Multi-slide clustering results on DLPFC. Clustering of tissue locations based on all 10 methods in the multi-slide clustering case across all 12 samples.

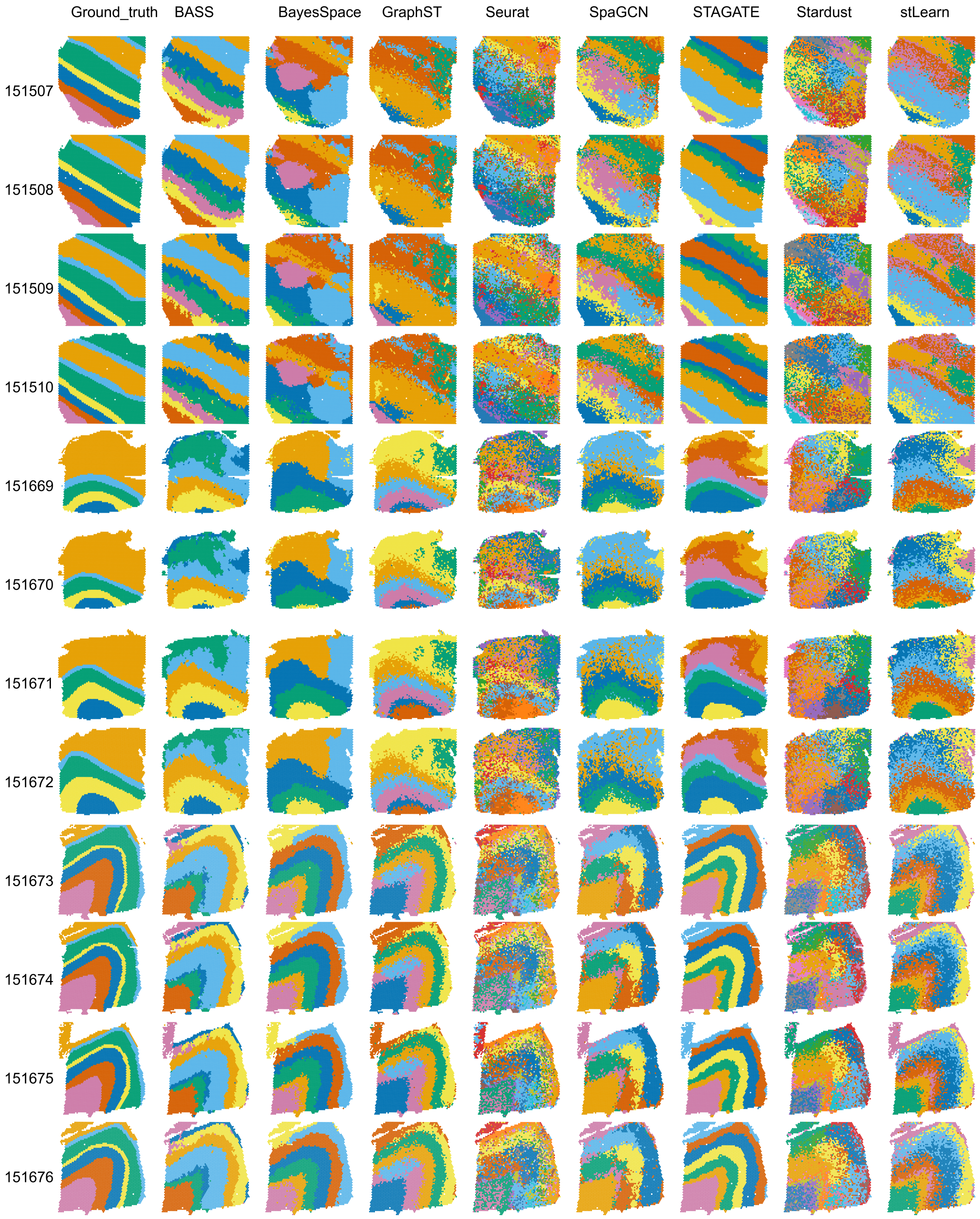


**Supplementary Figure 5.** Multi-slide clustering applied with harmony on DLPFC. Clustering of tissue locations based on BASS, BayesSpace, GraphST, Seurat, SpaGCN, STAGATE, Stardust, and stLearn in the multi-sample clustering applied with harmony across all 12 slides.

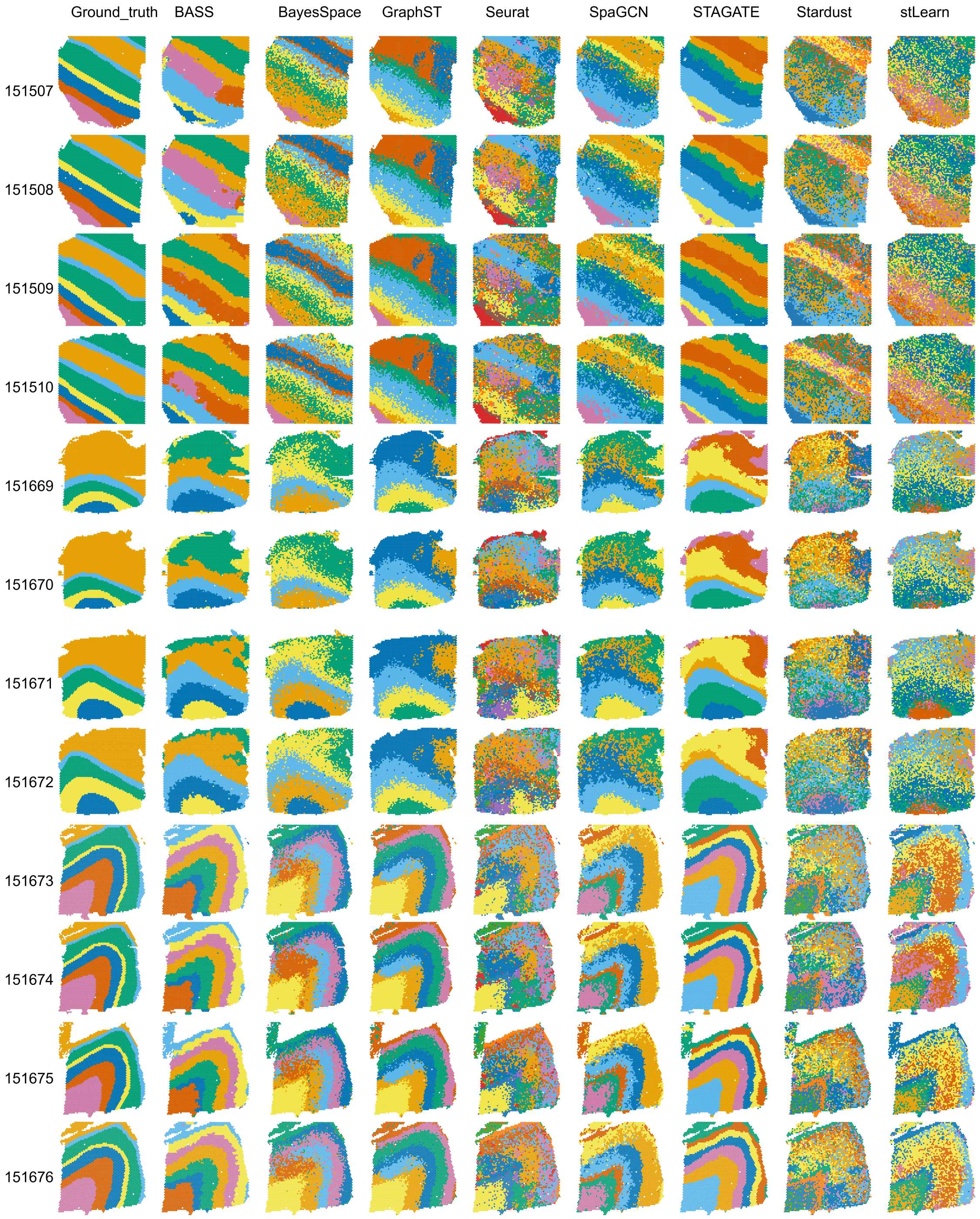


**Supplementary Figure 6.** Multi-slide clustering applied with PASTE on DLPFC. Clustering of tissue locations based on BayesSpace, GraphST, Seurat, SpaGCN, STAGATE, Stardust, and stLearn in the multi-slide clustering applied with PASTE across all 12 slides.

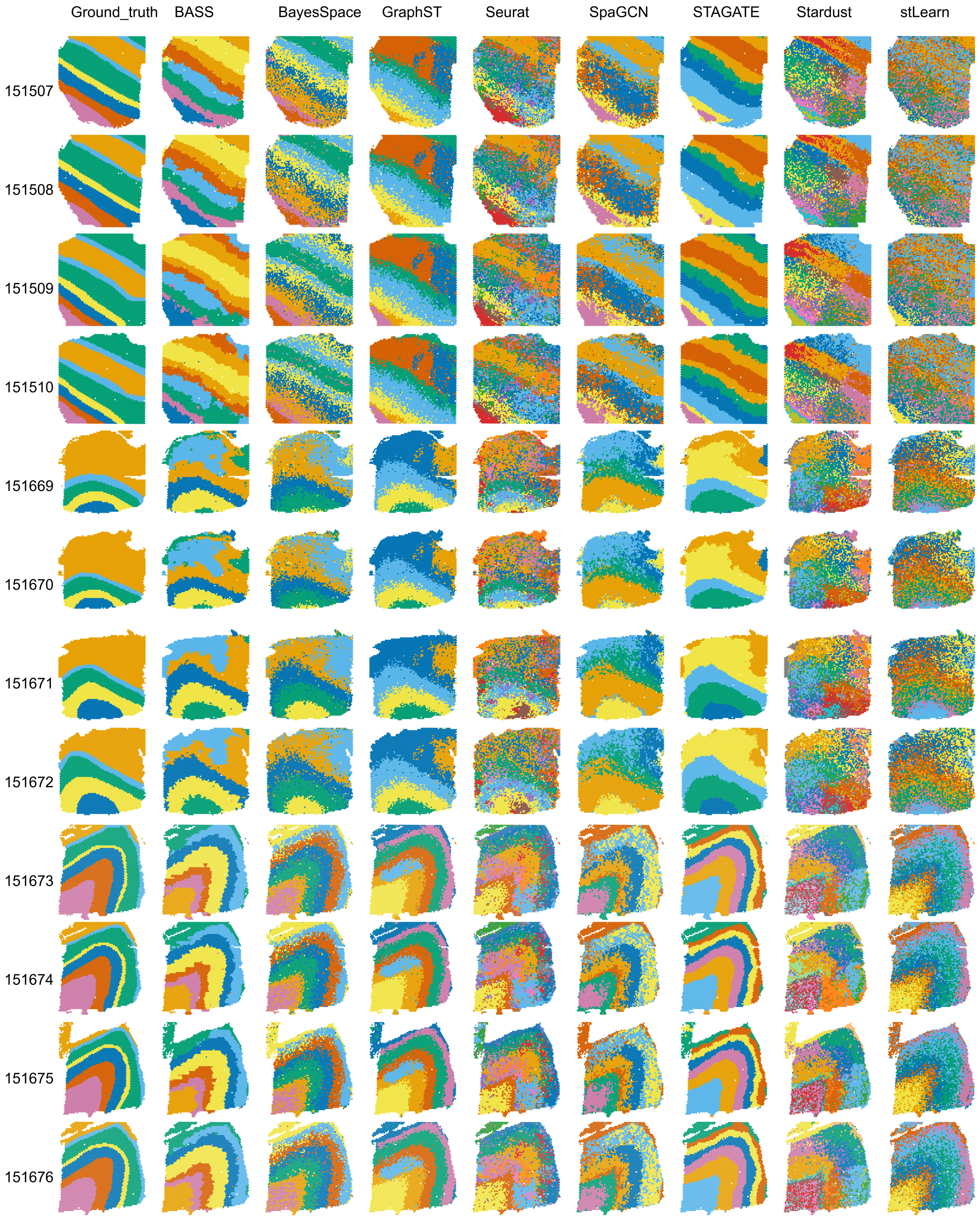


**Supplementary Figure 7.** Multi-slide clustering applied with both PASTE and Harmony on DLPFC. Clustering of tissue locations based on BayesSpace, GraphST, Seurat, SpaGCN, STAGATE, Stardust, and stLearn in the multi-slide clustering applied with both PASTE and Harmony across all 12 slides.

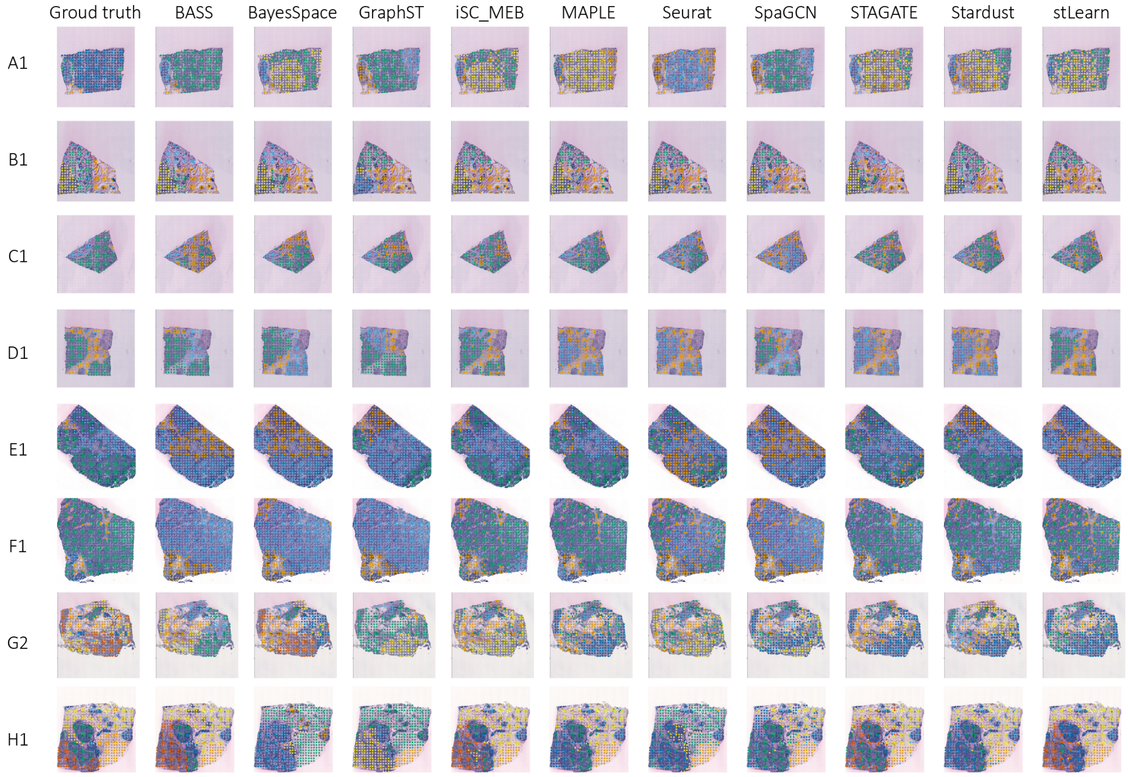


**Supplementary Figure 8.** Single-slide clustering results on HER2-positive breast cancer data. Clustering of tissue locations based on all 10 methods in the single-sample clustering case across 8 slides.

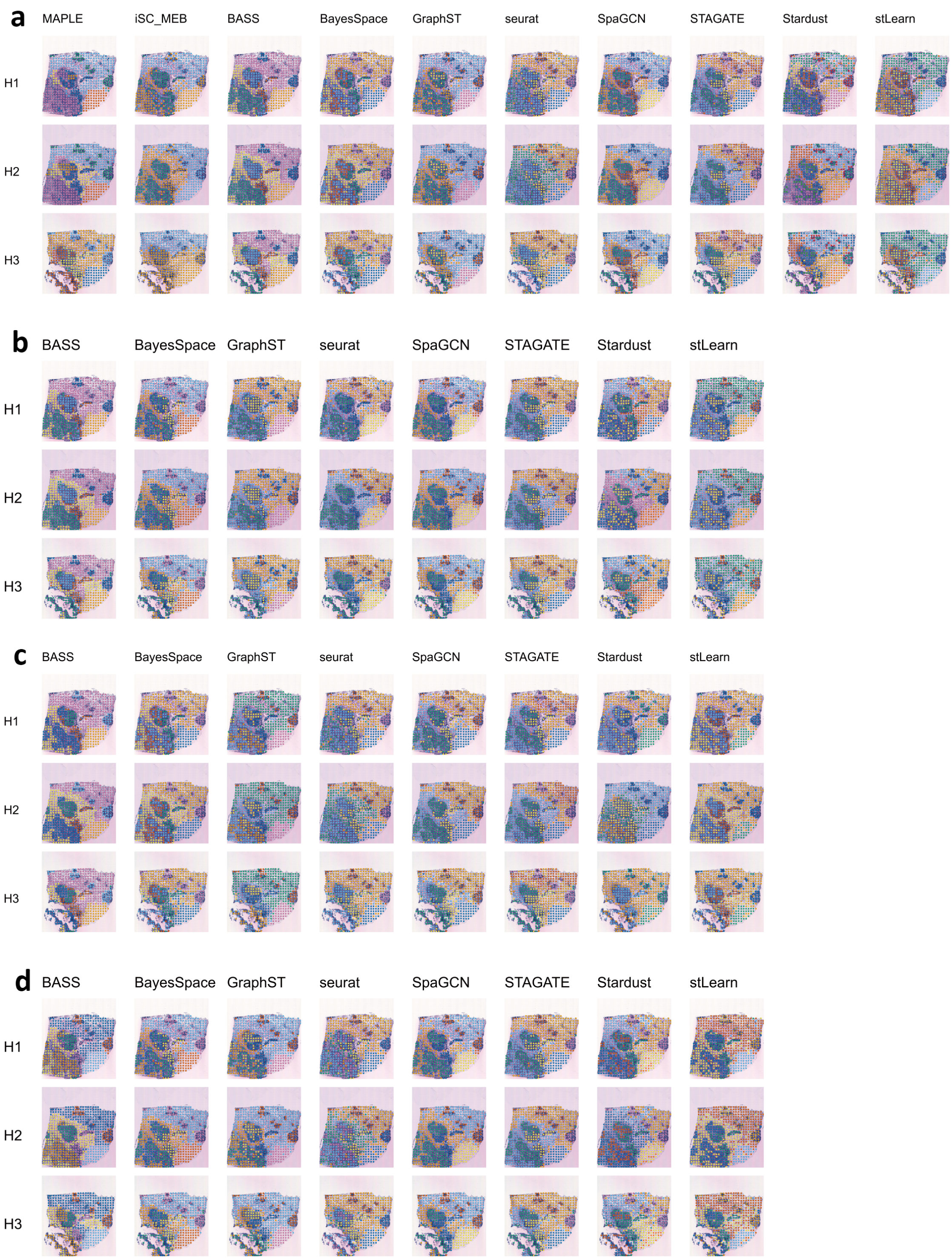


**Supplementary Figure 9.** Multi-slide clustering on HER2-positive breast cancer data H1-H3 with different techniques. **a.** Multi-slide clustering on H1-H3. **b.** Multi-slide clustering applied with Harmony on H1-H3. **c.** Multi-slide clustering applied with PASTE on H1-H3. d. Multi-slide clustering applied with PASTE and Harmony on H1-H3.

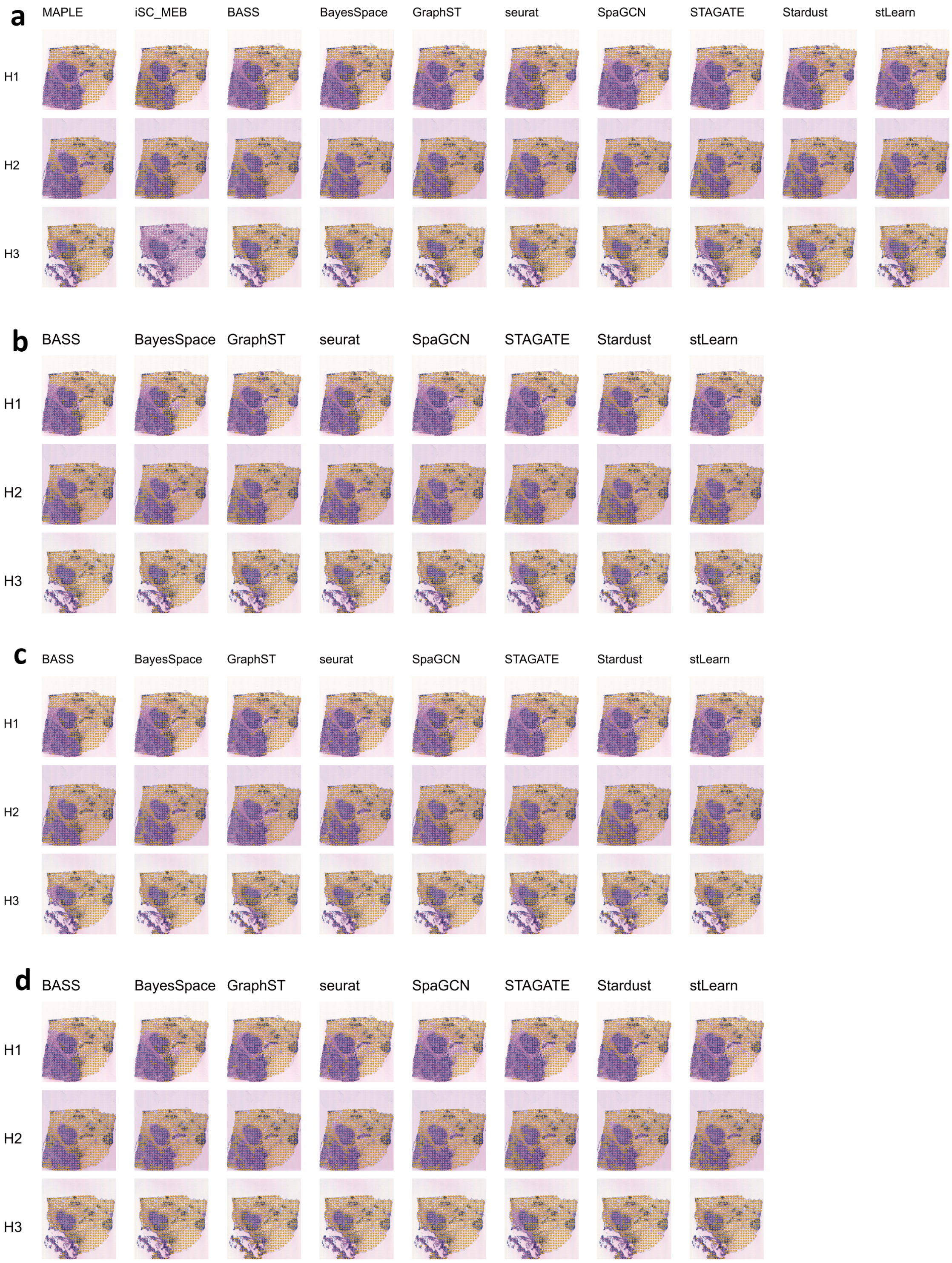


**Supplementary Figure 10.** Multi-slide clustering on HER2-positive breast cancer data H1-H3 with different techniques. **a.** The clustering results have been detected as cancer region and non-cancer region by cancer marker genes. Multi-slide Clustering on H1-H3. **b.** Multi-slide clustering applied with Harmony on H1-H3. **c.** Multi-slide clustering applied with PASTE on H1-H3. **d.** Multi-slide clustering applied with PASTE and Harmony on H1-H3.

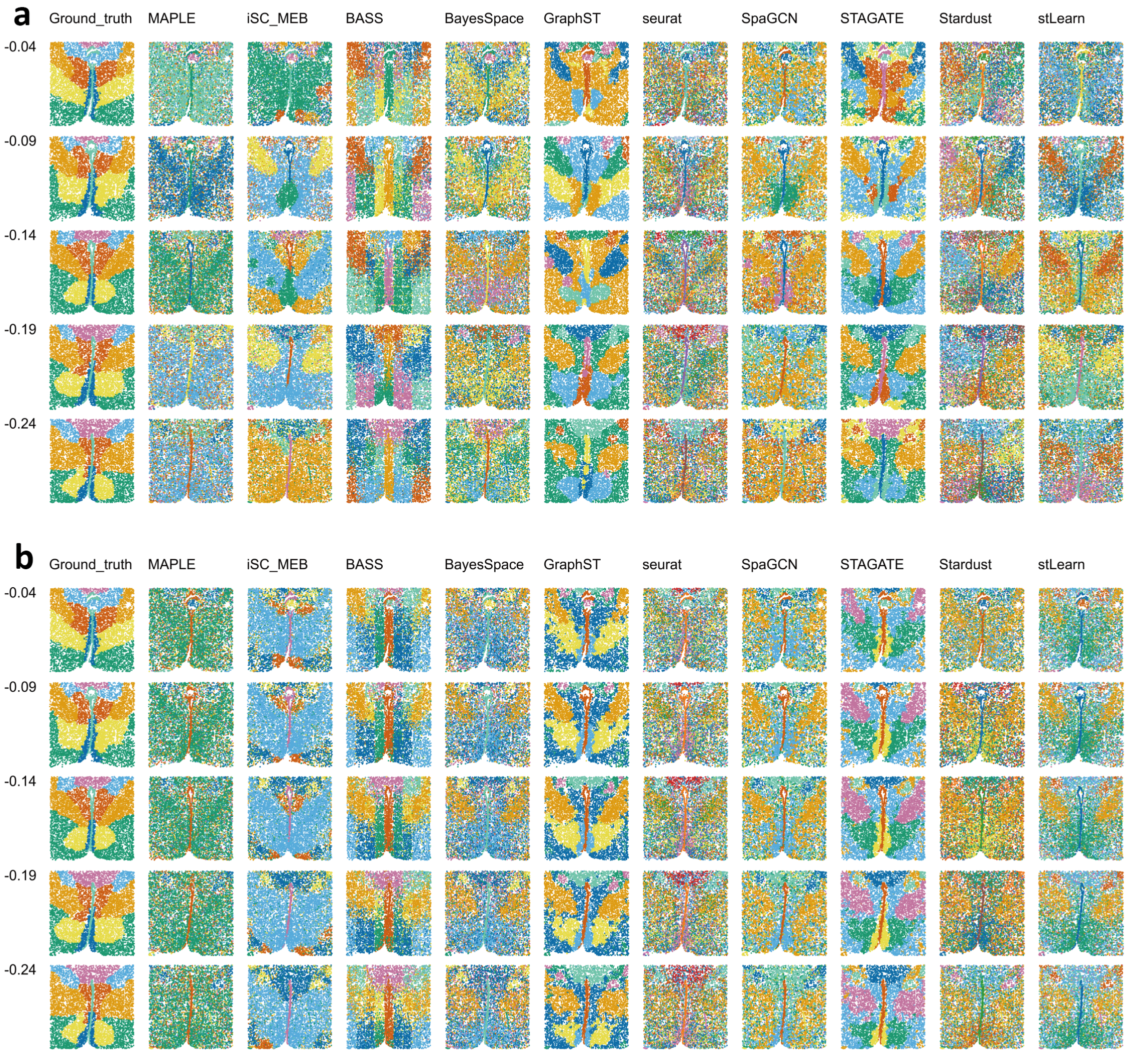


**Supplementary Figure 11.** 10 Clustering methods on mouse hypothalamus merfish data(Bregma -0.04,-0.09,-0.14,-0.19,-0.24). **a.** Single-slide clustering on 5 sections separately. **b.** Multi-slide clustering on 5 sections simultaneously.

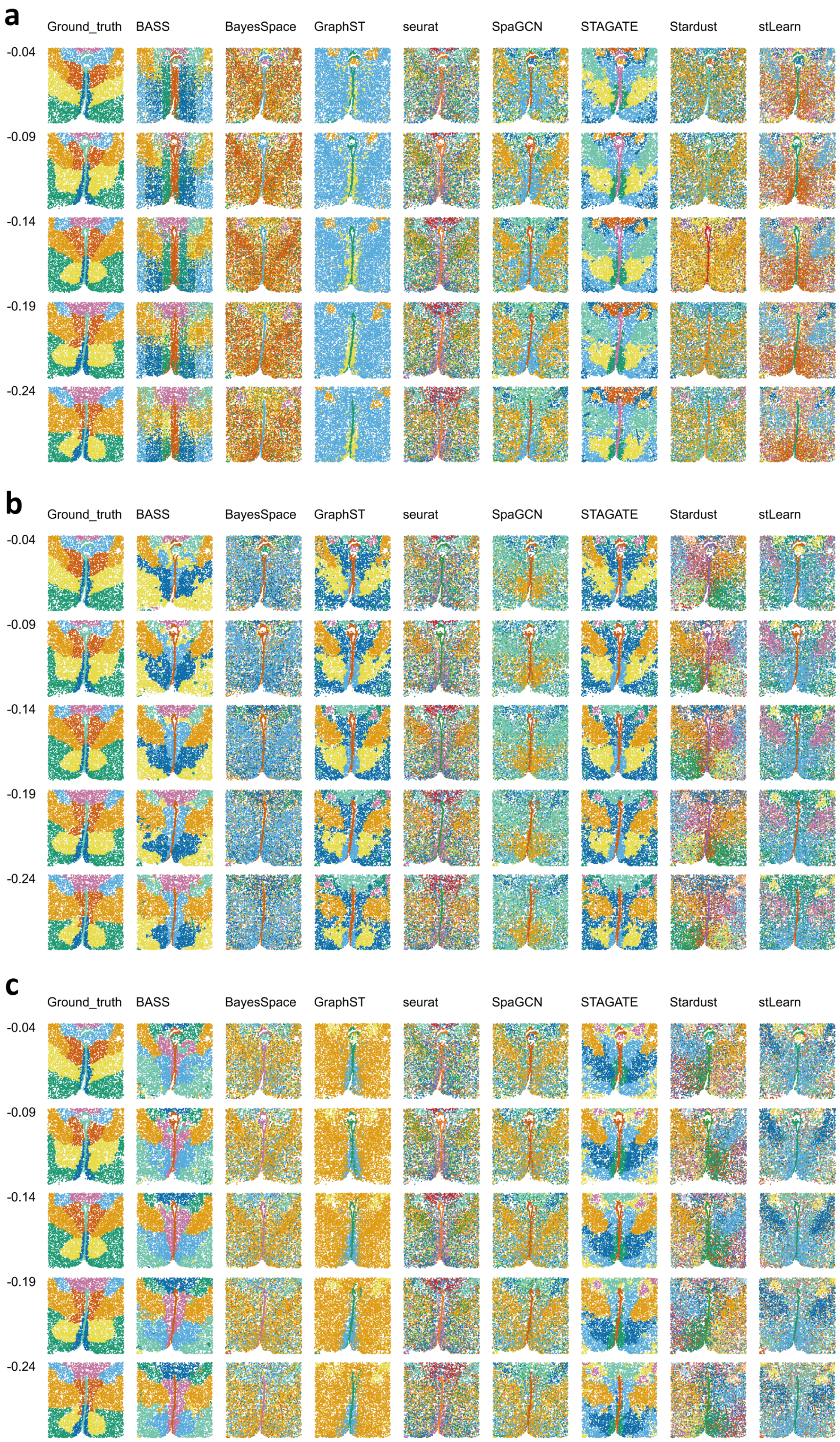


**Supplementary Figure 12.** 8 Clustering methods on mouse hypothalamus merfish data(Bregma -0.04,-0.09,-0.14,-0.19,-0.24). **a.** Multi-slide clustering applied with Harmony on 5 sections separately. **b.** Multi-slide clustering applied with PASTE on 5 sections simultaneously. **c.** Multi-slide clustering applied with PASTE and Harmony on 5 sections simultaneously.
